# Overexpression of the receptor-like kinase *BIR1* gene causes *SOBIR1*- and *EDS1*-dependent cell death phenotypes in Arabidopsis

**DOI:** 10.1101/2023.06.23.546234

**Authors:** Irene Guzmán-Benito, Carmen Robinson, Isabel Punzón, César Llave

## Abstract

The receptor-like kinase *BAK1-INTERACTING RECEPTOR-LIKE KINASE 1* (*BIR1*) functions as a negative regulator of multiple resistance signaling pathways in *Arabidopsis thaliana*. Previous studies showed that loss of *BIR1* function leads to extensive cell death and activation of constitutive immune responses. Here, we use a dexamethasone (DEX)-inducible expression system to investigate the impact of *BIR1* overexpression on plant growth and development and immune regulation. We show that, in the absence of microbes or microbe-derived elicitors/effectors, plants that overexpress *BIR1* displayed cell death phenotypes that concur with the transcriptomic up-regulation of multiple immune pathways involved in pathogen perception and defense signal transduction. *BIR1* overexpression has similar loss-of-function effects to *BIR1* depletion in knockout plants, which suggests that proper *BIR1* homeostasis requires a tight regulation of *BIR1* expression within a functional threshold. We show that *ENHANCED DISEASE SUSCEPTIBILITY 1* (*EDS1*) and *SUPPRESSOR OF BIR1-1* (*SOBIR1*) are required for the effector-triggered immunity (ETI)-type cell death phenotypes associated with overexpression of *BIR1*. Our data is then consistent with the current hypothesis by which loss of *BIR1* regulation and/or integrity is sensed by one or several guarding resistance (NLR) proteins to initiate a cell death response, in which SOBIR1 cooperates with EDS1 to transduce signals downstream of R proteins.

**Summary Statement:** Regulation of the receptor-like kinase *BIR1* has a strong impact on plant growth and development and immune homeostasis in Arabidopsis. *BIR1* overexpression causes cell death-and senescence-like phenotypes that require *EDS1* and *SOBIR1* signaling pathways, and that resemble to those observed by *BIR1* depletion.

## 1 INTRODUCTION

Plants use immunity to defend themselves against invaders such as bacteria, fungi, oomycetes, and viruses (Korner *et al*., 2013; Nishad *et al*., 2020). Pattern-triggered immunity (PTI) uses multiple cell-surface pattern recognition receptors (PRR) to recognize pathogen/microbe/damage-associated molecular patterns (PAMP/MAMP/DAMP). Effector-triggered immunity (ETI) involves intracellular receptors that sense pathogen-effector proteins or effector-induced manipulations of host proteins (Zhou & Zhang, 2020; Yu *et al*., 2021). PRRs are single-transmembrane receptor-like kinases (RLK) or receptor-like proteins (RLP). PRRs form active receptor complexes in a ligand-dependent manner with members of the somatic embryogenesis co-receptor kinases (SERKs), including BRASSINOSTEROID INSENSITIVE1-ASSOCIATED RECEPTOR KINASE 1 (BAK1/SERK3) (Boutrot & Zipfel, 2017; Hohmann *et al*., 2017; Smakowska-Luzan *et al*., 2018). BAK1 functions as a positive regulator of PTI (Wu *et al*., 2020), and has a dual role as positive and negative regulator of cell death during ETI (Belkhadir *et al*., 2012; Dominguez-Ferreras *et al*., 2015). RLP-type PRRs lack kinase activity and constitutively associate in a ligand-independent manner with the leucine-rich repeat (LRR)-RLK SUPPRESSOR OF BIR1-1 (SOBIR1) (Gust *et al*., 2017). Once the microbe is sensed, PRR complexes activate receptor-like cytoplasmic kinases (RLCKs) that initiate downstream signaling cascades (Liang & Zhou, 2018; Rao *et al*., 2018). Among them, BOTRYTIS INDUCED KINASE1 (BIK1) functions as a positive regulator of immunity when triggered by LRR-RLKs or as a negative regulator when triggered by LRR-RLPs (Bi *et al*., 2018; Wan *et al*., 2019).

ETI relies on nucleotide-binding domain LRR intracellular immune receptors (NLRs) encoded by polymorphic disease resistance genes. NLRs comprise either proteins with an N-terminal domain similar to the TOLL/INTERLEUKIN RECEPTOR 1 (TIR1) (TIR-NB-LRR, TNL), proteins with a coiled-coil (CC) domain at the N terminus (CC-NB-LRR, CNL) or proteins with a RESISTANCE TO POWDERY MILDEW 8-LIKE DOMAIN (RPW8)-type domain (RNLs) (van Wersch *et al*., 2020; Wang & Chai, 2020). NLRs function as “sensors” that recognize effectors, or as “helpers” that mediate signaling downstream of sensor NLRs (Jubic *et al*., 2019). Recent studies demonstrated that PTI and ETI work together to mutually potentiate the immune response (Yuan *et al*., 2021b; Yuan *et al*., 2023). NLR-mediated signaling is dependent on multiple PTI-associated components such as BAK1, SOBIR1, or BIK1 in a PTI-independent manner, while TNL-coding genes that are up-regulated during early PTI boost immune responses (Ngou *et al*., 2021; Tian *et al*., 2021; Yuan *et al*., 2021a). ENHANCED DISEASE SUSCEPTIBILITY (EDS1) and PHYTOALEXIN DEFICIENT 4 (PAD4), which are essential in TNL-mediated immunity, also play a key role in PTI signaling in response to different microbial elicitors (Pruitt *et al*., 2021b; Tian *et al*., 2021). Furthermore, PTI and ETI outputs usually converge, albeit with different amplitude and duration (Yuan *et al*., 2021b). Early immune responses include apoplastic reactive oxygen species (ROS) burst, cytosolic calcium (Ca^+2^) influx and activation of Ca^+2^-dependent protein kinases (CPK), and activation of mitogen-activated protein kinases (MAPK). This response is followed by the transcriptional activation of defense processes that include accumulation of defense phytohormones, regulation of cell wall and secretome composition alteration (Li *et al*., 2020).

Plants deploy a variety of mechanisms to avoid the detrimental effects of hyperactivated defenses (Couto & Zipfel, 2016; Withers & Dong, 2017; Mithoe & Menke, 2018). Among them, BAK1-INTERACTING RECEPTOR-LIKE KINASE (BIR) proteins play a key role in defense attenuation. The current model postulates that, in the absence of microbes, BIR proteins constitutively associate with BAK1, maintaining the PRRs in a resting state. During ligand perception, BAK1 dissociates from BIR and becomes accessible to complex with PRR receptors (Gao *et al*., 2009; Halter *et al*., 2014; Imkampe *et al*., 2017). The Arabidopsis genome contains four *BIR* homologs. *BIR1* negatively regulates cell death signaling pathways that depend on BAK1, SOBIR1, EDS1 and PAD4, and that likely involve components of the TNL-mediated ETI response (Gao *et al*., 2009; Schulze *et al*., 2022). Arabidopsis *BIR2* and *BIR3* have critical roles in repressing PRR-mediated PTI responses (Halter *et al*., 2014; Imkampe *et al*., 2017). Furthermore, the TNL protein CSA1 interacts physically with BIR3 and senses perturbances in the BIR3/BAK1 complex to induce cell death (Schulze *et al*., 2022). *BIR4* has so far not been characterized.

*BIR1* expression is transcriptionally activated by multiple microbial and viral pathogens in a salicylic acid (SA)-dependent manner (Guzman-Benito *et al*., 2019). Under normal conditions, the *BIR1* gene undergoes transcriptional negative regulation by RNA-directed DNA methylation whereas a potent site-directed post-transcriptional silencing dynamically reinforces the action of the epigenetic regulation when *BIR1* is transcriptionally activated (Guzman-Benito *et al*., 2019). RNA silencing is thus a critical mechanism for the maintenance of *BIR1* homeostasis. Indeed, we previously showed that overexpression of either a human influenza hemagglutinin (HA)- or a mCherry-tagged version of *BIR1* using a *Tobacco rattle virus* (TRV)-based expression system or a transgenic dexamethasone (DEX)-inducible system, respectively, produced a range of senescence and cell death phenotypes in Arabidopsis (Guzman-Benito *et al*., 2019). Whereas the autoimmune response in *bir1-1* knockout mutants has been extensively characterized (Gao *et al*., 2009; Liu *et al*., 2016), little is known about the molecular basis of the phenotypes associated to an excess of *BIR1*. In this work, we use a previously characterized DEX-inducible system in Arabidopsis to investigate the immune components and pathways responsible for the phenotypes associated with *BIR1* gene overexpression in the absence of microbes or microbe-derived elicitors/effectors. We found that senescence and cell death-like phenotypes in *BIR1* overexpressing lines coincide with an inadequate overexpression of genes encoding immune receptors and co-receptors involved in the perception and downstream signaling of ligands and effectors, which resemble those observed in *BIR1* knockout plants. We evaluated the effects of the genetic depletion of several master regulators of the immune response and found that the cell death phenotypes associated with *BIR1* overexpression require the TNL-associated EDS1 and/or the RLP-co-receptor SOBIR1, but the underlying mechanism is unknown.

## 2 METHODS

### 2.1 Plant material and growth condition

*Arabidopsis thaliana* plants were grown in controlled environmental chambers under long-day conditions (16h day/8h night) at 22°C. Arabidopsis lines used in this study were derived from the Columbia-0 (Col-0) ecotype. Seeds were surface-sterilized and sown on plates containing ½ Murashige and Skoog (MS) medium supplemented with MES (0,5g/L), 1% sucrose and 0,9% Plant agar (Duchefa Biochemie). Seedlings were transferred to soil 7 to 10 days after germination.

Mutants for *bir1-1* and *sobir1-12* were donated by Yuelin Zhang (University of British Columbia, Canada). The *eds1-2*, *pad4-1* and *eds5-3* alleles were supplied by Jane E. Parker (Max Planck Institute for Plant Breeding Research, Germany). Mutant *bak1-5* was donated by Birgit Kemmerling (Center of Plant Molecular Biology, University of Tübingen, Germany). The *sid2-2* mutant was a gift from Francisco Tenllado (Centro de Investigaciones Biológicas Margarita Salas, CSIC, Spain). All Arabidopsis plants used in this study were in the Col-0 background.

Arabidopsis transgenic plants expressing a BIR1-mCherry fusion under a dexamethasone (DEX)-inducible gene expression system (BIR1 L6 and BIR1 L9) were previously described (Guzman-Benito *et al*., 2019). The same DEX-inducible expression system containing a mCherry-tagged *BIR1* gene was transformed into Arabidopsis mutant backgrounds according to standard floral dipping (Clough & Bent, 1998). Independent homozygous lines were selected in T3 and used for subsequent experiments. Morphological phenotypes after DEX treatments were investigated in at least two independent transgenic lines of each genotype.

### 2.2 RNA and Protein analyses

Total RNA was extracted using TRIzol reagent (Invitrogen) and treated with DNase I (Invitrogen) following manufacturer’s instructions to eliminate genomic DNA traces. One-step quantitative RT-PCR (RT-qPCR) was carried out using Brilliant III Ultra-Fast SYBR Green QRT-PCR Master Mix (Agilent Technologies) in a Rotor-Gene 6000/Rotor-Gene Q real-time PCR machine (Corbett/Qiagen) (Fernandez-Calvino *et al*., 2016). Relative gene expression was determined using the Delta-delta cycle threshold method and Rotor-Gene 6000 Series Software (Corbett). Transcript levels of the target genes were normalized to the transcript levels of the housekeeping gene *CBP20* (*At5g44200*) as a reference gene. Gene-specific primers are given in Table S1.

For protein analyses, leaf tissue samples were ground in liquid nitrogen and homogenized in extraction buffer (65 mM Tris-HCl, pH 8; 3% sodium dodecyl sulfate [SDS]; 1% ß-mercaptoethanol; 10% glycerol). Samples were diluted in Laemmli buffer, incubated for 5 min at 95°C, and loaded onto 10% SDS-PAGE protein gels. After electrophoresis, proteins were blotted onto an ECL nitrocellulose membrane (Amersham-Pharmacia), and detected using commercial horseradish peroxide (HRP)-conjugated secondary antibodies and a chemiluminescent substrate (LiteAblot Plus).

### 2.3 Library preparation for transcriptome sequencing and bioinformatic analyses

Total RNA was extracted from each sample using the RNeasy plant mini kit (Qiagen) followed by DNase treatment according to manufacturer’s instructions. Each sample consisted of RNA pooled from six individual plants. The RNA purity and RNA concentration were checked using a Nanodrop spectrophotometer (ThermoFisher Scientific), while the RNA integrity was measured using the Agilent Bioanalyzer 2100 system (Agilent Technologies). Sequencing libraries were generated at NOVOGENE (HK) COMPANY LIMITED (Wan Chai, Hong Kong) (www.novogene.com) using NEBNext® UltraTM RNA Library Prep Kit for Illumina® (New England Biolabs) following manufacturer’s recommendations and index codes were added to attribute sequences to each sample. First-strand cDNA from mRNA was synthesized using random hexamer primer and M-MuLV Reverse Transcriptase (RNase H-). Second-strand cDNA synthesis was subsequently performed using DNA Polymerase I and RNase H. NEBNext Adaptors with hairpin loop structure were ligated. In order to select cDNA fragments of preferentially 150∼200 bp in length, the library fragments were purified with the AMPure XP system (Beckman Coulter). PCR was done with Phusion High-Fidelity DNA polymerase, Universal PCR primers and Index (X) Primer. PCR products were purified with the AMPure XP system and library quality was assessed on the Agilent Bioanalyzer 2100 system. Clustering of the index-coded samples was performed on a cBot Cluster Generation System using the TruSeq PE Cluster Kit v3-cBot-HS (Illumina) according to the manufacturer’s recommendations. Finally, paired-end reads were generated via an Illumina PE150 Hiseq platform at NOVOGENE (HK) COMPANY LIMITED.

Clean reads were obtained by removing adaptor tags and reads containing poly-N and low-quality reads from raw data. Paired-end clean reads were mapped to the Arabidopsis TAIR v10 reference genome using HISAT2 software. To quantify the gene expression levels, HTSeq v0.6.1 was used to calculate the number of mapped reads of each gene and then normalize the results to the expected number of Fragments Per Kilobase of transcript sequence per Millions of base-pairs sequenced (FPKM). Differential expression analysis between groups (three biological replicates per condition) was performed using DESeq2 R package (1.18.0). The resulting *p-value*s were adjusted using the Benjamini and Hochberg’s approach for controlling the False Discovery Rate (FDR). Genes with an adjusted *p-value* ≤0.05 found by DESeq2 were deemed as differentially expressed (DEG). Functional classification of DEGs including Gene Ontology (GO) and KEGG was performed. GO enrichment analysis of DEGs was implemented by tGOseq R package, in which gene length bias was corrected. KOBAS software was used to test the statistical enrichment of DEGs in the KEGG database GO terms and KEGG pathways with corrected *p-value* ≤0.05 were considered significantly enriched by DEGs. The raw data was deposited in the Gene Expression Omnibus under accession number GSE234036.

### 2.4 Dexamethasone (DEX) treatments and sampling procedure

DEX (Sigma) was dissolved in 100% ethanol to make a 30 mM DEX stock solution which was stored at –20°C in a light-tight vial. Arabidopsis plants were grown for three to four weeks in soil and sprayed at 24 h intervals with DEX at 30 µM as indicated (McNellis *et al*., 1998). At this concentration, plants display optimal induction of the transgene (Guzman-Benito *et al*., 2019). Samples for gene expression assays were collected at the outset of symptoms appearance, which normally corresponded to approximately 12 days of DEX treatments. This time however could differ between genotypes and treatments and consequently plants of each genotype may exhibit different ages and developmental stages when harvested compared to each other genotypes. As a result, quantitative comparisons and inferences on gene expression levels between genotypes should be made with caution in this study.

## 3 RESULTS

### 3.1 *BIR1* overexpression causes transcriptome reprogramming in Arabidopsis

We conducted a transcriptomic analysis of Arabidopsis BIR1 L9 plants, which express a DEX-inducible *BIR1-mCherry* transgene (hereafter *BIR1*) (Guzman-Benito *et al*., 2019), to understand the downstream biological processes and pathways associated with *BIR1* overexpression. This system proves to be a valuable tool for studying *BIR1* effects in the absence of pathogen elicitors or effectors. After DEX treatment, BIR1 L9 plants start showing morphological and growth defects from approximately day 12 onward, including severe stunting, leaf deformation, premature senescence (yellowing), and cell death (Figure 1a) (Guzman-Benito *et al*., 2019). At this time, we detected roughly two orders of magnitude more mCherry-tagged BIR1 protein than after four days of DEX treatment, indicating that BIR1 protein accumulates gradually overtime in DEX-treated BIR1 L9 plants (Figure 1a). These phenotypes were identical to those observed when a BIR1-HA-coding construct was systemically overexpressed in Arabidopsis plants using a TRV-based expression system (Figure 1a) (Guzman-Benito *et al*., 2019). Hence, we can confidently attribute the observed cell death phenotypes to overexpression of *BIR1* and not to DEX side-effects or protein tagging. Under our experimental conditions, and consistent with previous studies, DEX caused a moderate growth retardation in WT plants when applications began at an early growth stage (Boyes stage 1.5), whereas DEX caused no growth defects when applied at later stages of plant development (Boyes stage 1.7) (Figure 1a, S1a) (Boyes *et al*., 2001; Ouwerkerk *et al*., 2001). In no case, WT plants exhibited cell death or morphological phenotypes upon DEX treatment (Figure 1a, S1a).

**Figure 1.**
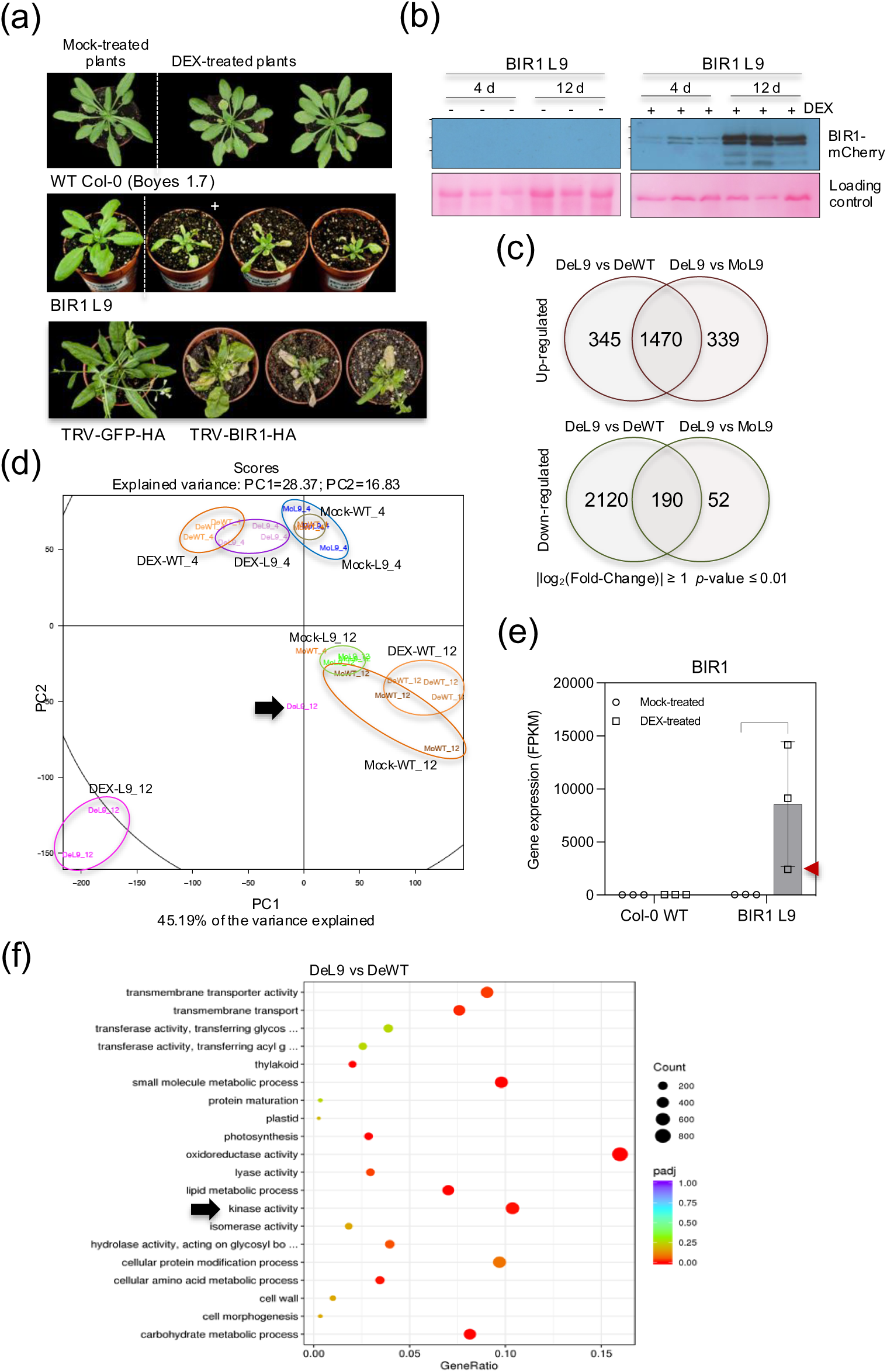
Transcriptional reprogramming associated with *BIR1* induction in Arabidopsis. **(a)** Morphology of representative dexamethasone (DEX)-inducible BIR1-mCherry transgenic Arabidopsis plants from line 9 (L9). Photographs were taken after 10-12 days of 30 µM DEX. Mock-treated wild-type (WT) Arabidopsis were used as controls. DEX treatments started at the Boyes growth stage 1.7. Cell-death phenotypes in Arabidopsis DEX-treated BIR1 L9 plants were similar to those observed when BIR1-HA was expressed via TRV infection (TRV-BIR1). **(b)** Western blot using anti-mCherry antibody shows the relative accumulation of BIR1-mCherry protein after 4 and 12 days of DEX treatments in BIR1 L9 plants. **(c)** Number of up-or down-regulated transcripts, with a |log_2_ (Fold-Change)| ≥1 and adjusted *p*-value ≤0.01, in each pairwise comparison is indicated. **(d)** Principal component analysis (PCA) of RNA-Seq data. The PCA was performed using normalized RNA-Seq data of differentially expressed genes (DEGs) under each condition: Mock-treated WT (Mock-WT), Mock-treated BIR1 L9 (Mock-L9), DEX-treated WT (DEX-WT), DEX-treated BIR1 L9 (DEX-L9). Samples were collected after 4 and 12 days of DEX treatment as indicated. Each biological replicate is represented in the score plot. The variance explained by each component (%) is given in parentheses. The sample (DeL9_12b) with differential expression within the DEX-treated L9 group is indicated with an arrow. **(e)** Relative expression of the BIR1-coding gene based on the number of Fragments Per Kilobase of transcript sequence per Millions of base-pairs sequenced (FPKM) (top). Significant differences between DEX- and mock-treated samples were analyzed using two-way ANOVA followed by Sidak’s multiple comparison test; ***, adjusted *p*-value < 0.001. **(f)** Scatterplot of selected gene ontology (GO) terms of DEGs. GO terms between WT and BIR1 L9 plants after 12 days of DEX-treatment (DeL9_ vs DeWT_) are shown. Dot size represents the count of different genes and the color indicates significance of the term enrichment. GO terms with adjusted *p*-value < 0.05 are significantly enriched. The kinase term (indicated by an arrow) was exclusively assigned to BIR1 L9 plants.

For RNA-seq, three-week-old plants were sprayed with DEX for 12 days whereas control plants were mock-treated with H_2_O. Three biological replicates were used as representative samples of each condition tested in this study: DEX-treated WT (DeWT_), mock-treated WT (MoWT_), DEX-treated BIR1 L9 (DeL9_) and mock-treated BIR1 L9 (MoL9_) plants as indicated (Table S2). To avoid extensive damage on the plant transcriptome, RNA seq samples were collected before the onset of cell death symptoms. Our data indicate that *BIR1* overexpression triggered a massive transcriptional reprogramming in Arabidopsis (Figure 1b) (Table S2). Differentially expressed genes (DEG) from different comparisons are presented in Table S3. DEGs using different statistical thresholds are shown in Figure S2a,b and Table S4.

Principal component analysis (PCA) was used to sort RNA-seq-based transcriptomic data according to gene expression levels. PC1 clearly separated the DEX-treated BIR1 L9 samples (DeL9_12) from the other conditions in the PCA score plot after 12 days of treatment, indicating that the *BIR1* expression level was the main source of variance (28.37%) (Figure 1c). WT samples, irrespective of DEX treatment, and mock-treated BIR1 L9 samples clustered in their proximity, reflecting similar gene expression profiles (Figure 1c). This result also suggested that DEX alone was not a major contributor of the transcriptomic changes observed in DEX-treated BIR1 L9 samples. PC2 highlights a shift in gene expression between samples due to age or developmental stage (16.83%) (Figure 1c). Samples of both genotypes collected after 4 days of DEX treatments were located close each other in the score plot, suggesting that, under this induction conditions, *BIR1* has a minimal impact on gene expression. Intriguingly, the DeL9_12b sample was located far from the other DeL9_12 samples in the PCA plot, and near the samples within the DeWT_12, MoWT_12 or MoL9_12 groups (Figure 1c). The levels of *BIR1* induction in the replicate DeL9_12b was substantially lower than in the other two replicates within the DeL9_12 group (Figure 1d). This observation implies that a lower overexpression of *BIR1* in the DeL9_12b sample has much less impact on the plant transcriptome than a higher overexpression as shown for the other two DeL9_12 replicates.

Gene ontology (GO) analysis showed that genes encoding proteins with oxidoreductase activity, transmembrane transport and other metabolic processes or genes related to lipid or carbohydrate metabolism were particularly overrepresented in our whole set of DEGs, including WT and BIR1 L9 plants (Figure 1e, S2c) (Table S5). These alterations on the plant transcriptome were presumably DEX-dependent. In contrast, DEGs with kinase activity were only enriched in *BIR1* overexpressing plants, relative to mock-treated BIR1 L9 plants (Figure 1e and S2c) or DEX-treated WT plants (Figure S1c). The Kyoto Encyclopedia of Genes and Genomes (KEGG) pathway analysis showed the enrichment of genes related to several metabolic processes and genes involved in the biosynthesis of secondary metabolites that were specifically activated after DEX-treatments (Figure S3) (Table S6). Conversely, DEGs involved in plant hormone signal transduction were significantly enriched only in DEX-treated BIR1 L9, but not in mock-treated BIR1 L9 plants or DEX-treated WT plants.

### 3.2 *BIR1* overexpression triggers the expression of immunity-related genes

RNA-Seq results were further validated in mock- and DEX-treated BIR1 L9 plants after 12 days of treatment using RT-qPCR. We found that expression of DEGs *BIR1*, *PR1*, *RESPIRATORY BURST OXIDASE HOMOLOG PROTEIN D* (*RBOHD*), *FRK1* and *CYTOCHROME P450 MONOOXYGENASE 79B2* (*CYP79B2*) were all up-regulated in Arabidopsis plants overexpressing *BIR1* compared to mock-treated controls. This finding is consistent with their RNA-Seq-based profiles and confirms reproducibility in their expression patterns (Figure 2a).

**Figure 2.**
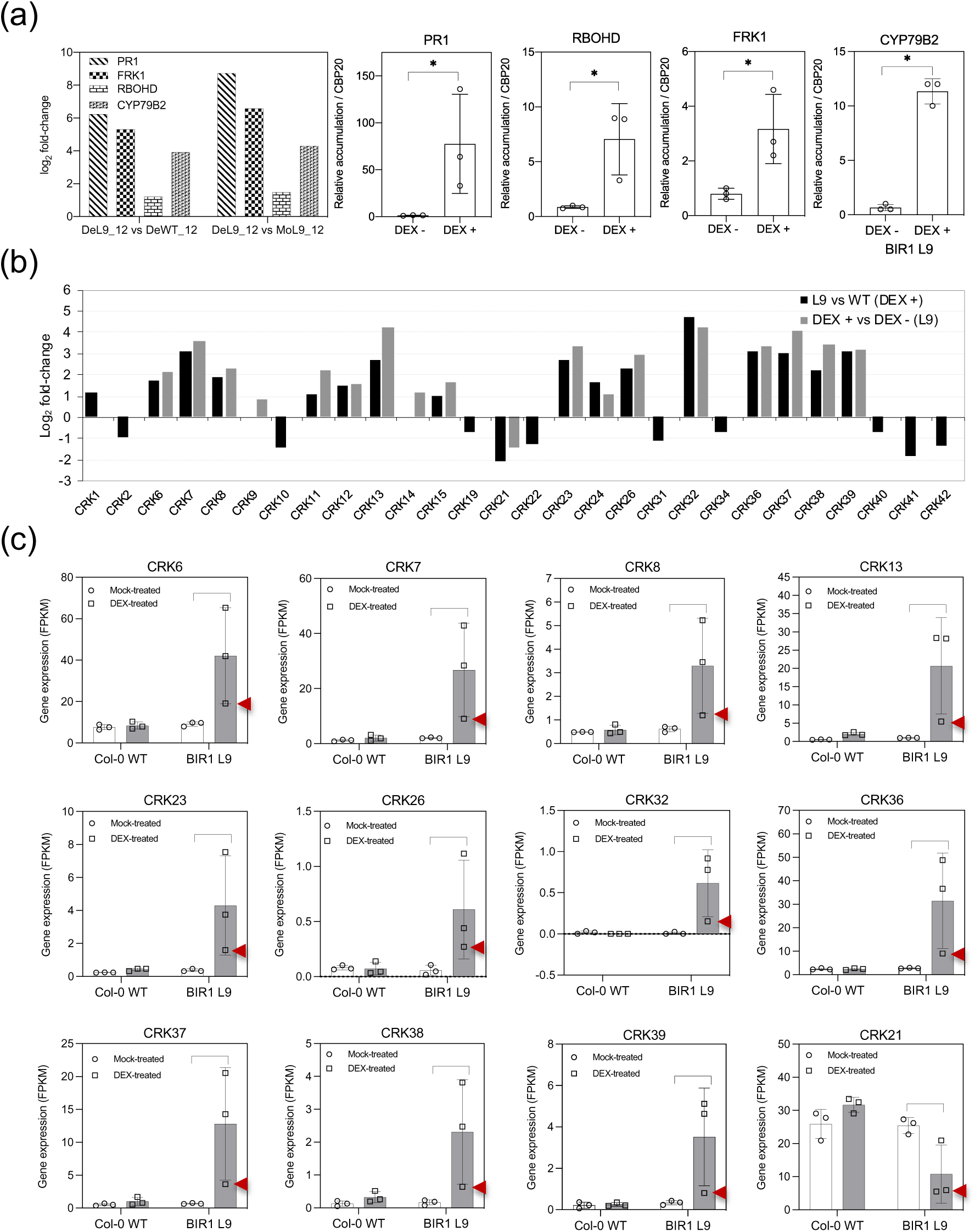
Differential expression of cysteine-rich receptor-like kinase (CRK)-coding genes in *BIR1* overexpressing plants. **(a)** Expression of selected BIR1-responsive differentially expressed genes (DEGs) based on fold-change values (log_2_ DeL9_12/DeWT_12 and log_2_ DeL9_12/MoL9_12 as indicated) was confirmed by RT-qPCR. Values from biologically independent samples were normalized to the *CBP20* internal control, and related to untreated (DEX-) plants, set to a value of 1. Data are mean ± SD analyzed by Mann-Whitney test; *, *p* <0.05. **(b)** Expression of CRK-coding differentially expressed genes based on log_2_ fold-change in BIR1 overexpressing plants as indicated. (**c)** Relative expression levels of CRK-coding genes represented by the number of Fragments Per Kilobase of transcript sequence per Millions of base-pairs sequenced (FPKM) in each of the three independent samples per condition used for RNA-Seq. Significant differences between dexamethasone (DEX)- and mock-treated samples were analyzed using two-way ANOVA followed by Sidak’s multiple comparison test; ***, adjusted *p*-value < 0.001, **< 0.01. Note that the expression levels of each individual replicate within the DEX-treated BIR1 L9_12 group mirror the abundance of *BIR1* transcripts in each replicate. Values from the DeL9_12b sample are indicated by an arrowhead.

A detailed inspection of the RNA-Seq data revealed that multiple genes involved in the perception of pathogens and the activation of defense pathways were misregulated when *BIR1* was overexpressed, but not in DEX-treated WT plants or mock-treated BIR1 L9 transgenic plants [|log_2_ (Fold-Change)| ≥1 and adjusted *p*-value ≤0.01] (Table S4). Two-tailed Fisher’s exact tests revealed that genes encoding LRR-type RLKs in the Arabidopsis genome were significantly enriched (Table S8). Particularly, genes encoding cysteine-rich RLKs (CRK), which likely are important in the perception of apoplastic ROS and in the activation of PTI and cell death, were overrepresented in the pool of DEGs (Figure 2b) (Table S7) (Chen, 2001; Yeh *et al*., 2015; Quezada *et al*., 2019; Castro *et al*., 2021). Interestingly, inspection of transcript abundance in single replicates indicated that the differential expression of CRK-coding genes was closely related to *BIR1* levels (Figure 2c). We also detected a high percentage of PRRs of the RLP class, not expected by chance, in the list of DEGs, being their relative transcript abundance consistent with *BIR1* expression levels (Figure 3a,b) (Table S7) (Wang *et al*., 2008; Lv *et al*., 2016). RLPs have been observed to be predominantly expressed in plants undergoing leaf senescence, supporting the idea that *BIR1* overexpression accelerates a senescence program (Wang *et al*., 2008). In Arabidopsis, *RLP23*, *RLP30*, and *RLP42* genes mediate SOBIR1-dependent immune activation, but the role of other RLPs in the perception of pathogen and defense is unknown (Wang *et al*., 2008; Zhang *et al*., 2013; Albert *et al*., 2019) (Table S7).

**Figure 3.**
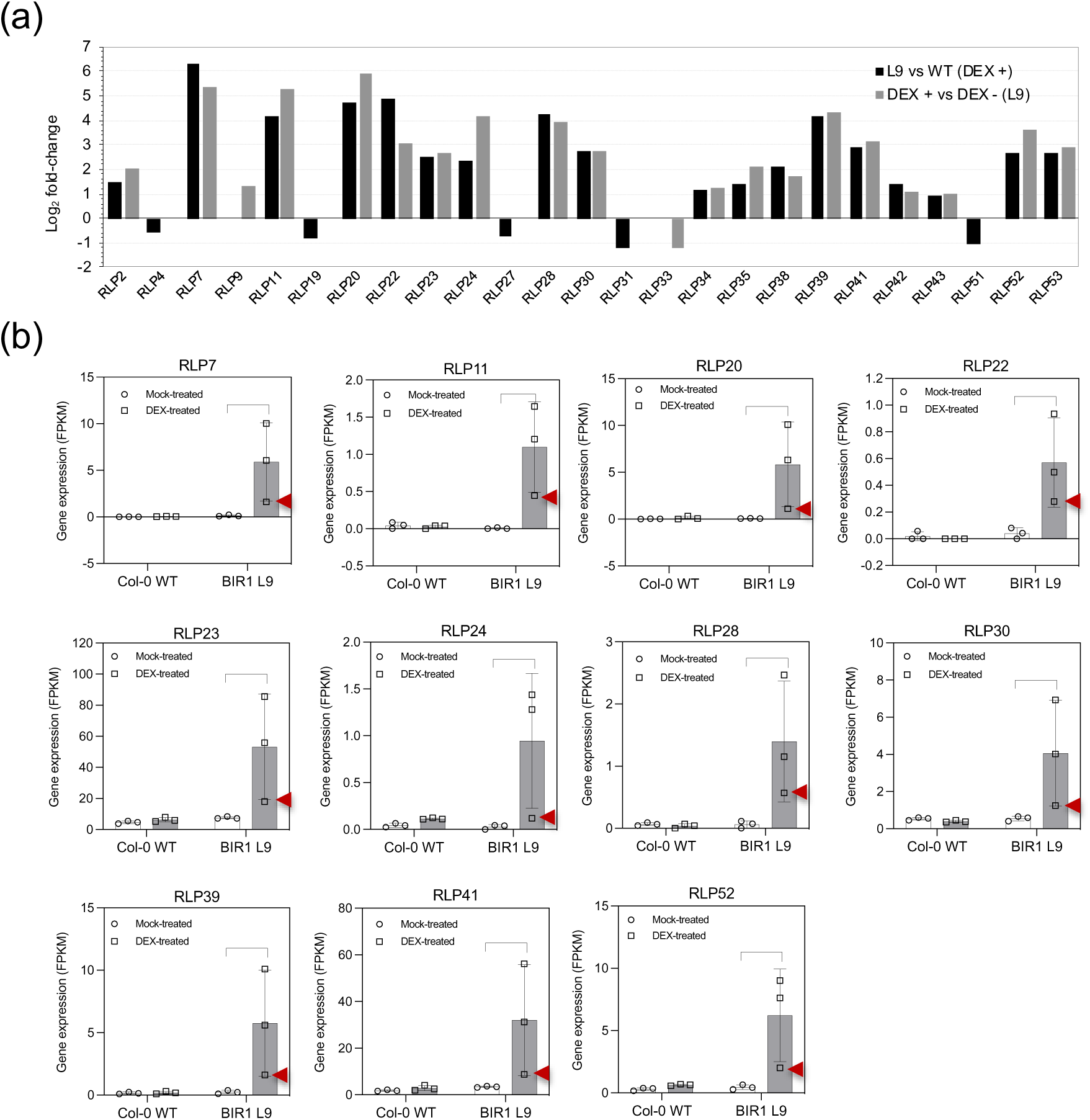
Differential expression of receptor-like protein (RLP)-coding genes in *BIR1* overexpressing plants. **(a)** Expression of differentially expressed RLP-coding genes based on log_2_ fold-change as indicated. **(b)** Relative expression levels of RLP-coding genes represented by the number of Fragments Per Kilobase of transcript sequence per Millions of base-pairs sequenced (FPKM) in each of the three independent samples per condition used for RNA-Seq. Significant differences between dexamethasone (DEX)- and mock-treated samples were analyzed using two-way ANOVA followed by Sidak’s multiple comparison test; ***, adjusted *p*-value < 0.001, **< 0.01. Note that the expression levels of each individual replicate within the DEX-treated BIR1 L9_12 group mirror the abundance of *BIR1* transcripts in each replicate. Values from the DeL9_12b sample are indicated by an arrowhead.

Conversely, PRRs of the RLK class known for eliciting immune responses were not collectively and significantly up-regulated in *BIR1* overexpressing plants (Figure S4a). For instance, the leucine-rich repeat RLK *ELONGATION FACTOR-TU RECEPTOR* (*EFR*) was up-regulated, whereas *FLAGELLING SENSING 2* (*FLS2*) or *PEP1 RECEPTOR 1* (*PEPR1*) and *PEPR2*, which sense danger-associated AtPeps (Krol *et al*., 2010), or *MDIS1-INTERACTING RECEPTOR LIKE KINASE 2* (*MIK2*), which recognizes conserved signature from phytocytokines and microbes (Hou *et al*., 2021), showed no responsiveness to *BIR1* overexpression. Among the lysin motif (LysM) RLK family (LYK), *LYK5* (Wan *et al*., 2012; Cao *et al*., 2014), or *LYSM DOMAIN GPI-ANCHORED PROTEIN 1* (*LYM1*), which perceives peptidoglycans from the bacterial cell wall (Willmann *et al*., 2011), were up-regulated in DEX-treated BIR1 L9 plants. In contrast, *LYK4* or *LYM2* displayed no responsiveness (Faulkner *et al*., 2013). The expression of the genes encoding lectin RLKs *LIPOOLIGOSACCHARIDE-SPECIFIC REDUCED ELICITATION* (*LORE*) (Luo *et al*., 2020) and *RESISTANT TO DFPM INHIBITION OF ABA SIGNALING 2* (*RDA2*) (Kato *et al*., 2022) was not affected by *BIR1* overexpression. Finally, we observed up-regulation of genes encoding the cell wall-associated kinases (WAK), *WAK1* and *WAK2*, which contain several extracytoplasmic epidermal growth factor (EGF) and recognize oligogalacturonides (He *et al*., 1996; Brutus *et al*., 2010).

In addition, *BIR1* overexpression boosted the transcription of many other genes involved in PTI and ETI signaling in a dose-dependent manner too (Figure 4a). This includes the RLK co-receptors *BAK1*, *SOBIR1* or *CERK1*, and RLCKs such as *BIK1* and several *AvrPphB SUSCEPTIBLE1(PBS1)-LIKE* (*PBL*) kinases (*PBL4*, *PBL12*, *PBL20*) (Liang & Zhou, 2018; Rao *et al*., 2018) (Table S7). Active BIK1 positively regulates the Ca^2+^ influx from the apoplast to the cytosol, which switches on the kinase activity of CPKs (Li *et al*., 2014; Yip Delormel & Boudsocq, 2019). Collectively, CPKs were not differentially regulated, but many genes encoding individual CPKs were regarded as DEGs in plants overexpressing *BIR1* (Table S3). For instance, the expression of *CPK4-6* and *CPK11* genes, which are key players in transcriptional reprogramming upon elicitor perception, was enhanced only in DEX-treated BIR1 L9 plants (Figure S5) (Boudsocq *et al*., 2010). Upon ligand perception, BIK1 dissociates from activated receptor complexes and phosphorylates the plasma membrane NADPH oxidase RBOHD to generate extracellular ROS (Kadota *et al*., 2014; Li *et al*., 2014). *RBOHD* was induced in DEX-treated BIR1 L9 plants, but not in their corresponding controls (Figure 4a). Genes encoding lipase-like proteins *EDS1*, *PAD4*, *SENESCENCE ASSOCIATED-GENE 101* (*SAG101*) and *ACTIVATED DISEASE RESISTANCE 1* (*ADR1*), which are required for TNL-receptor function during ETI, were also up-regulated (Figure 4a) (Table S3) (Zhou & Zhang, 2020; Ngou *et al*., 2021; Pruitt *et al*., 2021b; Sun *et al*., 2021; Tian *et al*., 2021). Interestingly, a large number of DEGs in our assay were annotated as resistance protein-coding genes, of which the TNL class was particularly enriched in *BIR1* overexpressing plants (Swiderski *et al*., 2009) (Table S7). Conversely, neither *NON-RACE SPECIFIC DISEASE RESISTANCE 1* (*NDR1*) nor genes encoding CNL intracellular receptors, which require *NDR1*, were differentially expressed in response to *BIR1* overexpression (Figure 4a) (Table S3). *EDS5,* an important regulator of SA biosynthesis in Arabidopsis (Nawrath *et al*., 2002), was up-regulated in DEX-treated BIR1 L9 samples.

**Figure 4.**
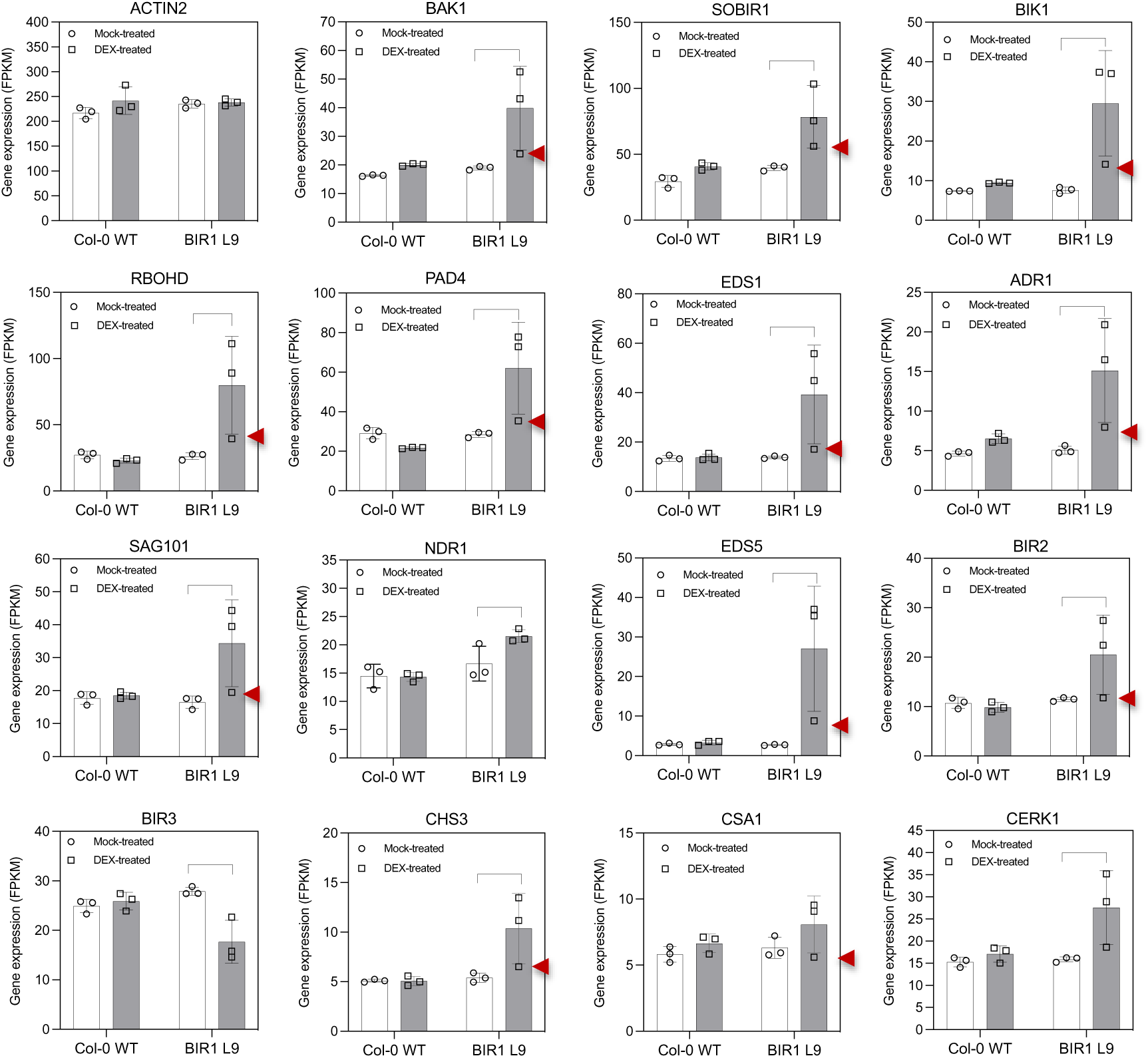
*BIR1* overexpression leads to the transcriptional activation of pattern-triggered immunity (PTI)-associated co-receptors and effector-triggered immunity (ETI) regulators. Relative expression of selected genes encoding PTI/ETI-markers based on the number of Fragments Per Kilobase of transcript sequence per Millions of base-pairs sequenced (FPKM) in each of the three independent samples per condition used for RNA-Seq. Actin expression is shown as control. Note that the expression levels of each individual replicate within the DEX-treated BIR1 L9_12 group mirror the abundance of *BIR1* transcripts in each replicate. DeL9_12b is indicated by a red arrow. Significant differences between mock-treated and DEX-treated samples were analyzed using two-way ANOVA followed by Sidak’s multiple comparison test; **, adjusted *p*-value < 0.01, * < 0.05. Values from the DeL9_12b sample are indicated by an arrowhead.

Interestingly, *BIR1* overexpression led to up-regulation of the *BIR2* gene, whereas *BIR3* and its partner *CONSTITUTIVE SHADE AVOIDANCE 1* (*CSA1*) genes were unaffected (Figure 4a) (Schulze *et al*., 2022). Finally, transcripts of plant heterotrimeric G proteins *ARABIDOPSIS G PROTEIN α-SUBUNIT1* (*GPA1*), *ARABIDOPSIS G PROTEIN β-SUBUNIT 1* (*AGB1*), *ARABIDOPSIS G PROTEIN γ-SUBUNIT 2* (*AGG2*), and *EXTRA-LARGE G PROTEIN 1* and *2* (*XLG1/2*) were more abundant in plants overexpressing *BIR1* relative to controls after 12 days of DEX treatment (Figure S6). Proteins encoded by these genes positively regulate stomatal defense and PTI signaling, ROS production and BIK1 stability allowing optimal immune activation (Liang *et al*., 2016; Zhong *et al*., 2019). Differential expression of immune-related genes occurred in *BIR1* overexpressing plants but not in DEX-treated WT plants, which discard a potential masking effect of DEX on defense gene expression. Consistent with transcript levels, SOBIR1, PAD4-dependent PR1 and EDS1 proteins were strongly up-regulated by *BIR1* overexpression in DEX-treated BIR1 L9 plants, compared to the non-treated controls (Figure S4b).

In conclusion, in the absence of microbes or microbe-derived elicitors/effectors, *BIR1* overexpression drastically interferes with the immune homeostasis, causing an inadequate overexpression of genes encoding immune receptors, particularly RLP and TNL receptors, and co-receptors involved in the perception and downstream signaling of ligands and effectors. We propose that the cell death-like phenotypes caused by *BIR1* overexpression, which are reminiscent of those observed when *BIR1* function is lost, are due to an intrinsic deregulation of multiple defense and resistance pathways.

### 3.3 Genetic disruption of *SOBIR1*, but not *BAK1*, partially rescues *BIR1* overexpression phenotypes

To understand the genetic basis of the phenotypes associated to *BIR1* overexpression, in the following sections, we examined if mutations in key components of PTI and ETI signaling alleviate the cell death-type morphologies and up-regulation of immune genes (Rodriguez *et al*., 2016; Freh *et al*., 2022). The DEX-inducible BIR1-mCherry-coding cassette was transformed into the different mutant backgrounds tested in this study. To overcome potential insertion position effects, two independent BIR1-mCherry transgenic lines of each mutant genotype were monitored. Phenotypes resulting from *BIR1* overexpression described in this work were consistently reproduced for each mutant in both independent transgenic lines and between experiments (Figure S7a,b). None of the morphological and growth defects observed in transgenic BIR1 L9 overexpressing *BIR1* occurred in the mutant controls (without the inducible BIR1-mCherry construct) or in a transgenic line with a mCherry-coding transgene after DEX treatments (Figure S8). These controls further discard that the observed phenotypes were caused by DEX treatments and/or mCherry overexpression (Figure S8).

*SOBIR1* and *BAK1* are necessary for activating cell death and constitutive defense responses in the *bir1-1* mutant (Gao *et al*., 2009; Liu *et al*., 2016). To test if they were also required for the *BIR1* phenotypes, we transformed the DEX-inducible BIR1-mCherry cassette into the *sobir1-12* and *bak1-5* mutants (Gao *et al*., 2009; Schwessinger *et al*., 2011). Plants were sprayed with DEX to trigger *BIR1* overexpression, or H_2_O as control. Western blot analysis showed that *sobir1-12* BIR1 and *bak1-5* BIR1 mutants accumulated as much BIR1-mCherry protein as BIR1 L9 plants after 7 days of DEX treatments, indicating similar expression of the *BIR1* transgene (Figure 5a). Consistent with previous results, BIR1 L9 plants, used as reference set throughout the experiment, displayed severe stunting, accelerated senescence (yellowing) and cell death, and accumulated abnormally high levels of *BIR1* transcripts after 12 days of DEX treatment (Figure 5a,c). Localized cell death in the *BIR1* overexpressing inducible system was previously shown using trypan blue staining (Guzman-Benito *et al*., 2019). When *BIR1* was overexpressed in the *sobir1-12* mutant, we observed milder phenotypes than those in BIR1 L9 plants, indicating that the *sobir1-12* mutation partially suppresses the morphological defects associated to *BIR1* overexpression (Figure 5b, S7a,b). *BIR1* transcripts levels in these same *sobir1-12* BIR1 plants remained strongly elevated (Figure 5c, S7c). The increased accumulation of transcripts of the PTI-markers *RBOHD*, *FRK1* and *WRKY DNA-BINDING PROTEIN 29* (*WRKY29*) observed upon *BIR1* overexpression was also drastically reduced in DEX-treated *sobir1-1*2 BIR1 plants (Figure 5d). These findings indicate that *BIR1* overexpression activates immune signaling pathways that require *SOBIR1*. *bak1-*5 BIR1 transgenic plants showed strong morphological phenotypes and severe yellowing after 12 days of DEX application (Figure 5b, S7a,b). In the set of i*bak1-*5 BIR1 plants shown in Figure 5b, *BIR1* transcript levels were only moderate (Figure 5c), but remained as high as in BIR1 L9 plants in other experimental repetitions (Figure S7c). *RBOHD*, *FRK1* and *WRKY* genes remained transcriptionally up-regulated in DEX-treated *bak1-5* BIR1 plants although at varying degrees (Figure 5e). Collectively, these data indicate that the morphological phenotypes and, perhaps, PTI gene activation, observed when *BIR1* is overexpressed are genetically unrelated to *BAK1*.

**Figure 5.**
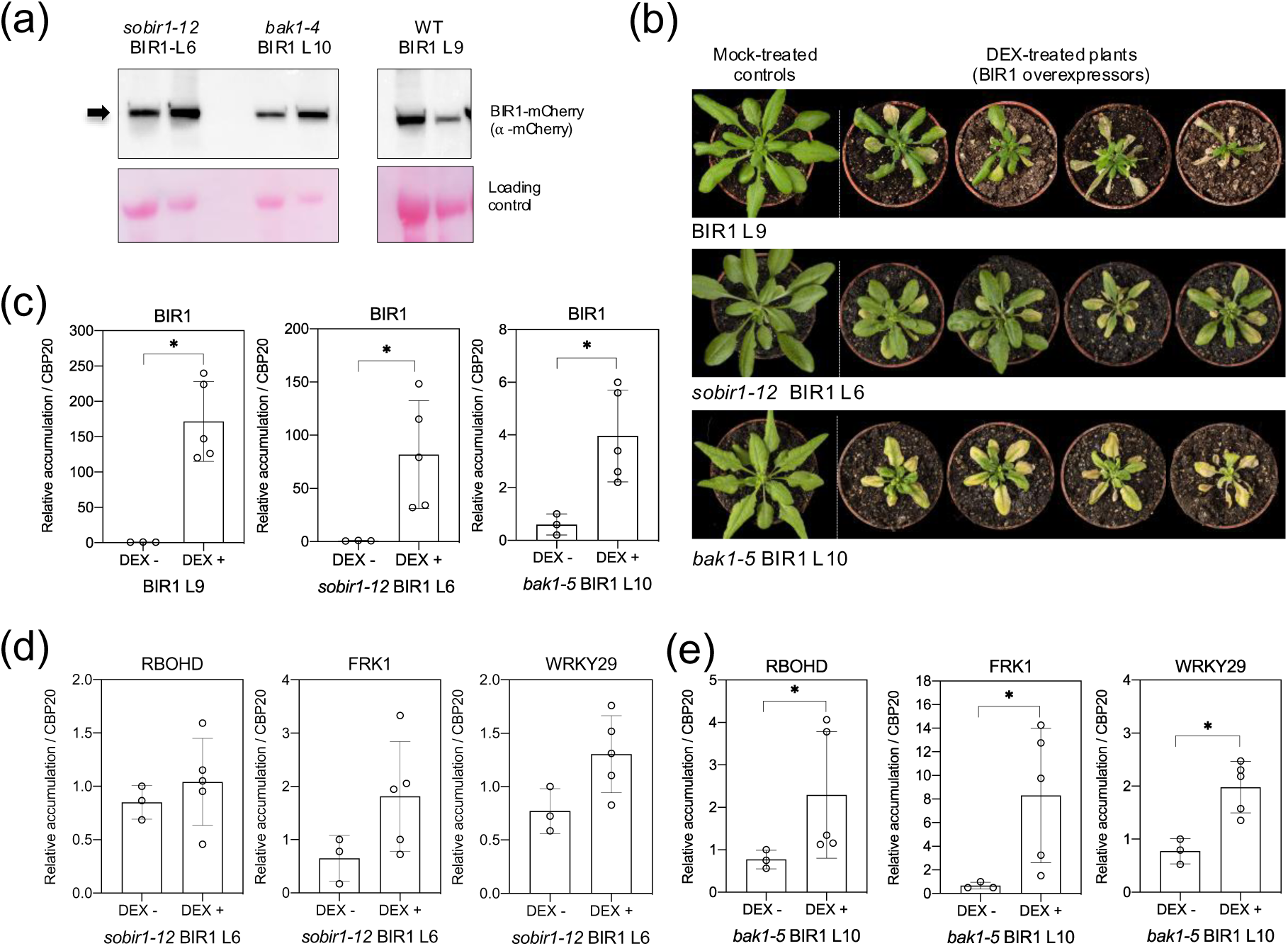
Phenotypes caused by *BIR1* overexpression are partially rescued in *sobir1-12* mutants. **(a)** Western blot analysis of BIR1-mCherry proteins in the dexamethasone (DEX)-inducible BIR1-mCherry transgenic Arabidopsis lines wild-type (WT) BIR1 L9, *sobir1-12* BIR1 (L6) and *bak1-* 5 BIR1 (L10) treated with DEX for 7 days. Each sample is a pool of two plants, and two samples per genotype are shown. Protein loading control by Ponceau S is shown. **(b)** Morphology of representative WT BIR1 L9, *sobir1-12* BIR1 L6 and *bak1-*5 BIR1 L10 plants. Images from independent transgenic lines are shown in **Figure S8**. **(c)** RT-qPCR analysis of *BIR1* transcripts in WT BIR1 L9, *sobir1-12* BIR1 L6 and *bak1-*5 BIR1 L10 plants. **(d, e)** RT-qPCR analysis of PTI-markers *RBOHD*, *FRK1* or *WRKY29* in *sobir1-12* BIR1 L6 **(d)** and *bak1-*5 BIR1 L10 **(e)** plants. Photographs and samples for qRT-PCR were taken after 12 days of 30 µM DEX treatment. Measures were done in mock-treated (DEX-) versus DEX-treated (DEX+) using the same set of replicates from each genotype. Values from biologically independent samples were normalized to the *CBP20* internal control, and related to mock-treated plants, set to a value of 1. Data are mean ± SD analyzed by Mann-Whitney test; *, *p* <0.05. Experiments were repeated at least twice with similar results.

### 3.4 Genetic disruption of *EDS1*, but not *PAD4*, partially rescues *BIR1* overexpression phenotypes

A previous report showed that stunting, activation of cell death and defense responses in the *bir1-1* mutant are partially dependent on *EDS1* and *PAD4* (Gao *et al*., 2009). The EDS1-PAD4 node mediates both PTI and ETI through PRR signaling and TNL activation (Pruitt *et al*., 2021a). To test if *BIR1* overexpression phenotypes were dependent on *EDS1* or *PAD4*, the DEX-inducible BIR1-mCherry system was used to generate homozygous transgenic lines from *eds1-2* and *pad4-1* mutants (Parker *et al*., 1996; Zhou *et al*., 1998). The resulting *eds1-2* BIR1 and *pad4-1* BIR1 plants produced comparable amounts of BIR1-mCherry protein to the BIR1 L9 control after 7 days of DEX applications (Figure 6a). After 12 days of DEX treatment, the *eds1-2* BIR1 plants displayed mostly WT-like appearance and, occasionally, localized yellowing after DEX, indicating that *eds1-2* can partially rescue the *BIR1* overexpression morphologies (Figure 6b, S7a). RT-qPCR analysis showed elevated levels of *BIR1* transcripts in these DEX-treated *eds1-2* BIR1 plants (Figure 6c). Since TNL-coding transcripts were abundant in *BIR1* overexpressing plants, we speculated that *EDS1*-dependent TNL-mediated resistance responses are activated when *BIR1* is overexpressed. We next found that the expression of the PTI-markers *FRK1* and *RBOHD* remained elevated in DEX-treated *eds1-2* BIR1 plants compared to non-treated controls (Figure 6d). *WRKY29* showed a tendency, albeit not significant, to be up-regulated in these plants (Figure 6d). Thus, our data indicated that the lack of *EDS1* attenuates the morphological phenotypes caused by *BIR1* overexpression, but has no obvious effects on PTI gene expression. DEX-inducible expression of *BIR1* in *pad4-1* BIR1 mutant plants produced growth defects similar to those in BIR1 L9 plants, albeit yellowing was less severe, indicating that mutations in *PAD4* were unable to rescue the *BIR1* overexpression phenotypes (Figure 6b,c, S7a). Interestingly, the PTI-components *FRK1*, *WRKY29* and *RBOHD* showed no responsiveness to *BIR1* overexpression in DEX-treated *pad4-1* BIR1 plants, which suggests that PTI gene activation in these plants requires *PAD4* (Figure 6e). In summary, the genetic elimination of *EDS1* mitigates the phenotypes caused by overexpression of *BIR1*, whereas induction of PTI gene expression in *BIR1* overexpressing plants seems to depend on *PAD4*.

**Figure 6.**
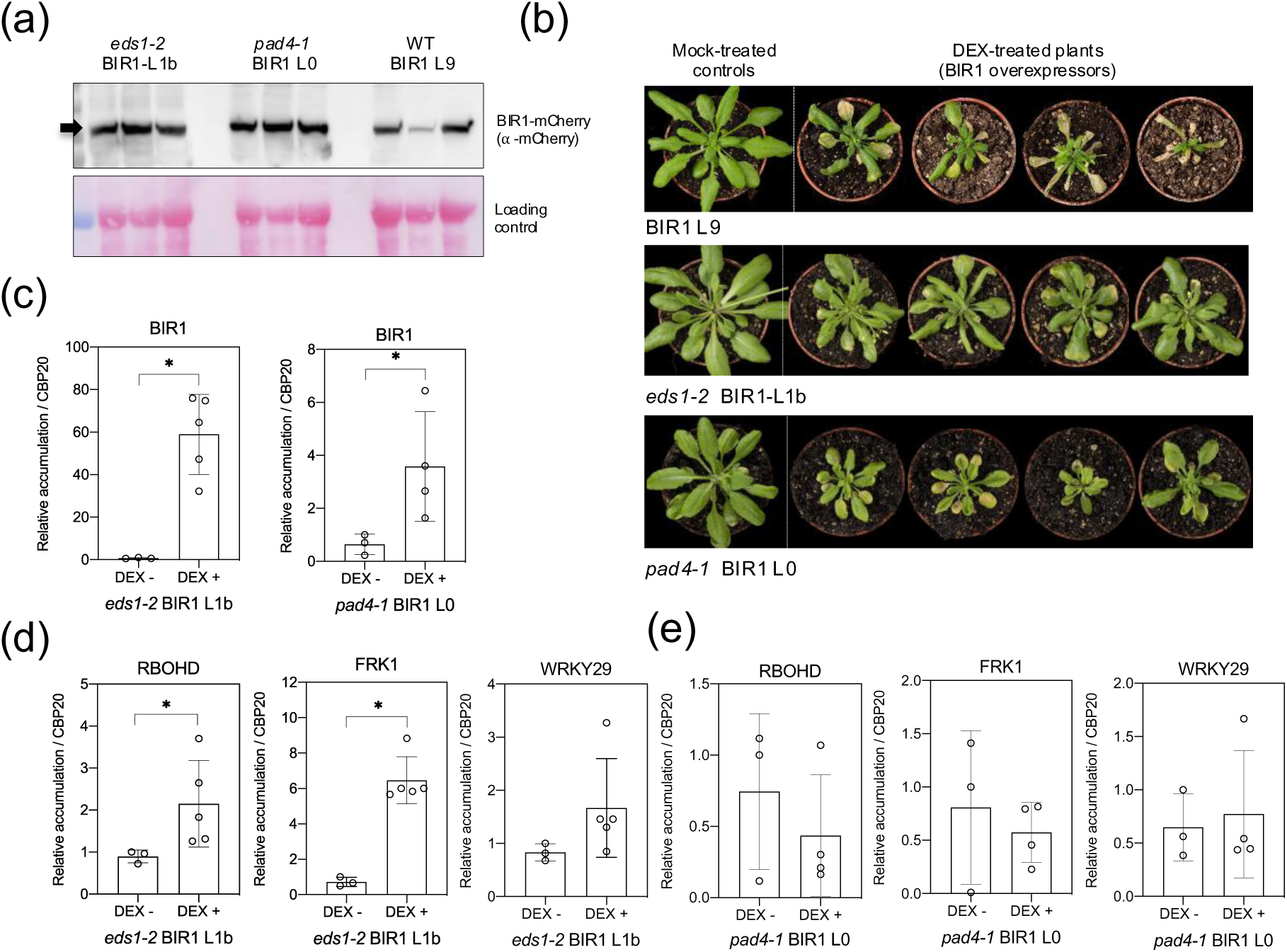
Phenotypes caused by *BIR1* overexpression are partially rescued in *eds1-2* mutants, but not in *pad4-1* mutants. **(a)** Western blot analysis of BIR1-mCherry proteins in the dexamethasone (DEX)-inducible BIR1-mCherry transgenic Arabidopsis lines wild-type (WT) BIR1 L9, *eds1-2* BIR1 L1b and *pad4-1* BIR1 L0 treated with DEX for 7 days. Each sample is a pool of two plants, and three samples per genotype are shown. Protein loading control by Ponceau S is shown. **(b)** Morphology of representative *eds1-2* BIR1 L1b and *pad4-1* BIR1 L0 plants. Images from independent transgenic lines are shown in Figure S8. **(c)** RT-qPCR analysis of *BIR1* transcripts in *eds1-2* BIR1 L1b and *pad4-1* BIR1 L0 mutant lines. **(d, e)** RT-qPCR analysis of PTI-markers *RBOHD*, *FRK1* or *WRKY29* in *eds1-2* BIR1 L1b **(d)** and *pad4-1* BIR1 L0 **(e)** plants. Photographs and samples for qRT-PCR were taken after 12 days of 30 µM DEX treatment. Measures were done in mock-treated (DEX-) versus DEX-treated (DEX+) using the same set of replicates from each genotype. Values from biologically independent samples were normalized to the *CBP20* internal control, and related to mock-treated plants, set to a value of 1. Data are mean ± SD analyzed by Mann-Whitney test; *, *p* <0.05. Experiments were repeated at least twice with similar results.

### 3.5 Phenotypes of *BIR1* overexpressing lines are unrelated to SA signaling pathways

Previously, high levels of SA were found to be partly responsible for *bir1-1* phenotypes (Gao *et al*., 2009). Here we tested the contribution of SA to the *BIR1* overexpression phenotypes. First, we used Arabidopsis deficient in the *SALICYLIC ACID INDUCTION DEFICIENT 2* (*SID2*)/ *ISOCHORISMATE SYNTHASE 1* (*ICS1*) and *EDS5* genes, which encode key SA biosynthetic enzymes (Huang *et al*., 2020), to infer their effects on the cell death phenotypes. DEX-inducible BIR1-mCherry transgenic plants were generated in the *sid2-2* and *eds5-3* (*eds5* is allelic to *sid1*) mutants, which contain low levels of SA (Nawrath *et al*., 2002). The resulting *eds5-3* BIR1 plants accumulated BIR1-mCherry protein like the BIR1 L9 control, whereas *sid2-2* BIR1 produced less protein after 7 days of DEX applications (Figure 7a). Nevertheless, *sid2-2* BIR1 and *eds5-3* BIR1 plants exhibited severe morphological phenotypes similar to those observed in BIR1 L9 plants, and also accumulated high levels of *BIR1* transcripts after 12 days of DEX treatment (Figure 7b, S7a). Thus, the *BIR1* overexpression phenotypes were likely unrelated to SA levels.

**Figure 7.**
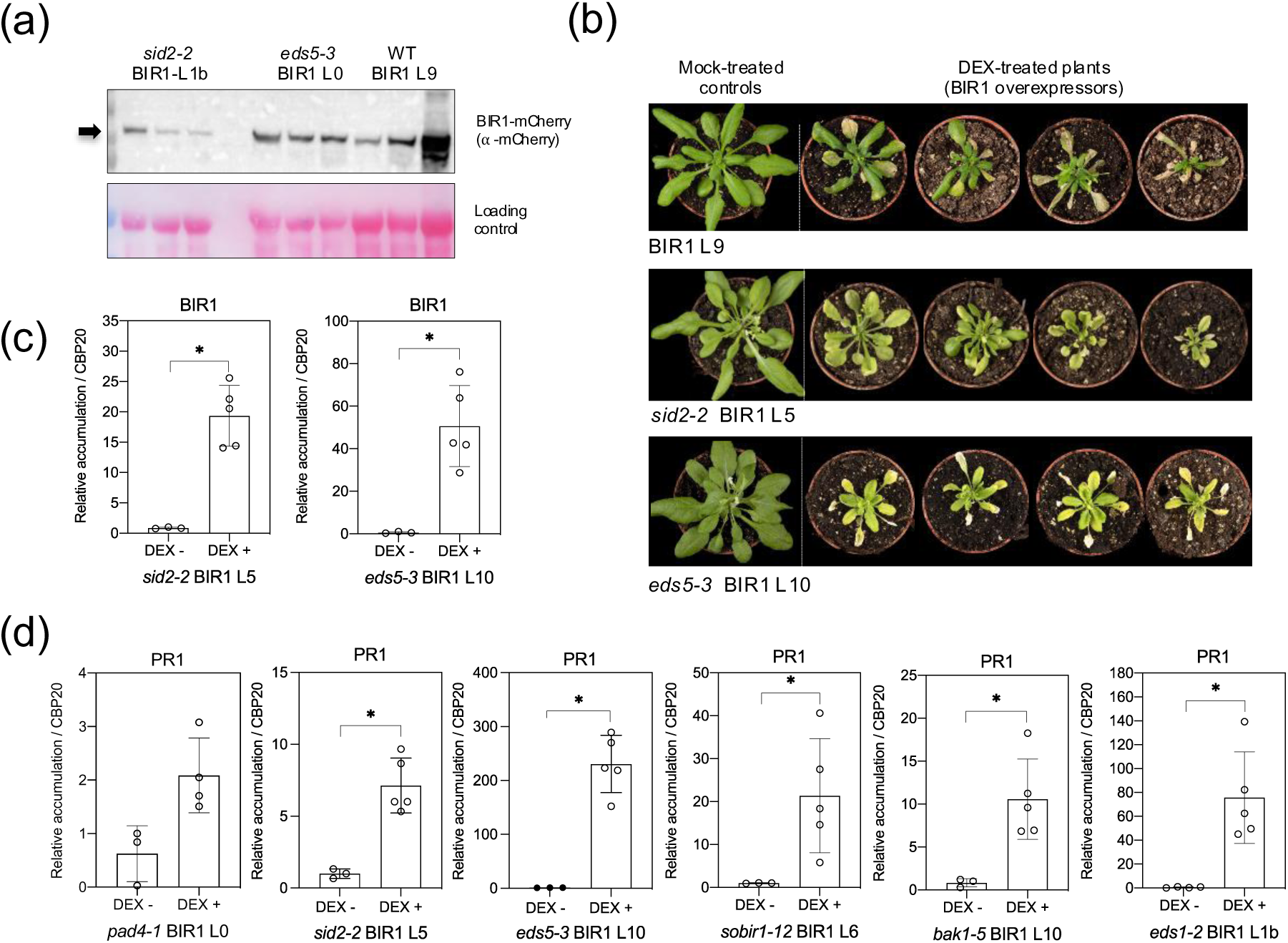
Phenotypes caused by *BIR1* overexpression are independent of salicylic acid (SA)-mediated signaling pathways. **(a)** Western blot analysis of BIR1-mCherry proteins in the dexamethasone (DEX)-inducible BIR1-mCherry transgenic Arabidopsis lines wild-type (WT) BIR1 L9, *sid2-2* BIR1 L5 and *eds5-3* BIR1 L10 plants treated with DEX for 7 days. Each sample is a pool of two plants, and three samples per genotype are shown. Protein loading control by Ponceau S is shown. **(b)** Morphology of representative *sid2-2* BIR1 L5 and *eds5-3* BIR1 L10 plants. Images from independent transgenic lines are shown in Figure S8. **(c)** RT-qPCR analysis of *BIR1* transcripts in *sid2-2* BIR1 L5 and *eds5-3* BIR1 L10 mutant lines. **(d)** RT-qPCR analysis of PR1 transcripts in *eds1-2* BIR1 L1b, *pad4-1* BIR1 L0, *sobir1-12* BIR1 L6, *bak1-*5 BIR1 L10, *sid2-2* BIR1 L5 and *eds5-3* BIR1 L10 mutant lines. Photographs and samples for qRT-PCR were taken after 12 days of 30 µM DEX treatment Measures were done in mock-treated (DEX-) versus DEX-treated (DEX+) using the same set of replicates from each genotype. Values from biologically independent samples were normalized to the *CBP20* internal control, and related to mock-treated plants, set to a value of 1. Data are mean ± SD analyzed by Mann-Whitney test; *, *p* <0.05. Experiments were repeated at least twice with similar results.

Interestingly, RT-qPCR revealed that transcripts of the canonical SA-related defense marker gene *PR1* were highly abundant in *BIR1* overexpressing *sid2-2* BIR1 *eds5-3* BIR1 plants compared to mock-treated controls (Figure 7d). The induction of *PR1* transcripts was also significant in SA-deficient, DEX-treated *sid2-2* BIR1 plants, but much less than in DEX-treated BIR1 L9 plants or *eds5-3* BIR1 plants (Figure 7d). Thus, our finding suggests that SA-independent mechanisms are also important for *PR1* expression in *BIR1* overexpressing plants. PR1 gene expression was also up-regulated in DEX-treated *sobir1-12* BIR1 and *bak1-5* BIR1 plants, irrespective of their different morphological phenotypes (Figure 7d). During immunity, *EDS1* and *PAD4* regulate SA-related defense gene expression and pathogen resistance through ICS1-generated SA-dependent and SA-independent mechanisms (Tsuda *et al*., 2013; Cui *et al*., 2017). In our study, *PR1* expression was strongly enhanced in *eds1-2* BIR1 plants, but not in *pad4-1* BIR1 plants, indicating that *PAD4*, but not *EDS1*, was required for *PR1* expression in *BIR1* overexpressing plants (Jirage *et al*., 1999) (Figure 7d). Together, our data propose that the sustained expression of *BIR1* activates both SA-dependent and SA-independent mechanisms of *PR1* induction whose contribution to *PR1* levels may differ between mutants.

### 3.6 Expression of phytohormone defense markers genes in compromised-defense mutants overexpressing *BIR1*

RNA seq data indicated a significant transcriptional reprogramming of genes involved in phytohormone metabolism and signaling in plants overexpressing *BIR1*. We determine the expression of several well-characterized phytohormone-related marker genes in the above Arabidopsis mutants defective in immune responses to infer the impact of plant hormone signaling on the defects observed in *BIR1* overexpressing plants. Transcript levels of jasmonic acid (JA)-responsive *PLANT DEFENSIN 1.2* (*PDF1.2*), ethylene ET- and JA-responsive *PR4* as well as genes encoding the cytochrome P450 enzymes CYP71B15 (*PAD3*) and CYP79B2 involved in camalexin (CA)-biosynthesis and CA-mediated plant defense were analyzed by RT-qPCR. None of these genes were affected by DEX in WT Arabidopsis plants.

Transcription of the *PDF1.2* gene was significantly up-regulated in all mutants tested, albeit to varying degrees (Figure 8a). Compared to other *BIR1* overexpressing mutants, *PDF1.2* transcripts were lower in *bak1-5* mutants, suggesting a role for *BAK1* in the regulation of JA-signaling in *BIR1* overexpressing plants, as previously reported for JA-dependent resistance against wounding or herbivory attacks (Figure 8a) (Yang *et al*., 2011; Tungadi *et al*., 2021). *EDS1* and, to a lesser extent, *PAD4* act as repressors of JA/ET defense signaling (Brodersen *et al*., 2006). Consistently, we found that induction of *PDF1.2* was strong in *eds1-2* BIR1 mutants, but weak, albeit significant, in *pad4-1* BIR1 plants (Figure 8a). The ET/JA marker *PR4* gene was poorly activated in all *BIR1* overexpressing mutants tested, except in the SA-defective *sid2-2* and *eds5-3* mutants, suggesting that the mutually antagonistic crosstalk between SA and ET/JA pathways is unaffected by *BIR1* overexpression (Stroud *et al*., 2022) (Figure 8b). The expression of CA-related *PAD3* and *CYP79B2* genes was up-regulated, albeit to variable extents, in all defense-compromised mutants tested upon *BIR1* overexpression, except *PAD4*-dependent *PAD3* which, as predicted, remained as the non-treated controls in *pad4-1* mutants (Zhou *et al*., 1999) (Figure 8c). Interestingly, the transcriptional activation of genes encoding markers in the JA, ET and CA pathways was particularly strong in DEX-treated *eds1-2* mutants, which display partially rescued phenotypes, but weak in *pad4-1* mutants, which display typical *BIR1* overexpression phenotypes. In summary, qRT-PCR analyses of canonical marker genes suggest that the cell death-like morphological phenotypes observed in *BIR1* overexpressing plants were not necessarily linked to the activation/inhibition of hormone defense signaling pathways.

**Figure 8.**
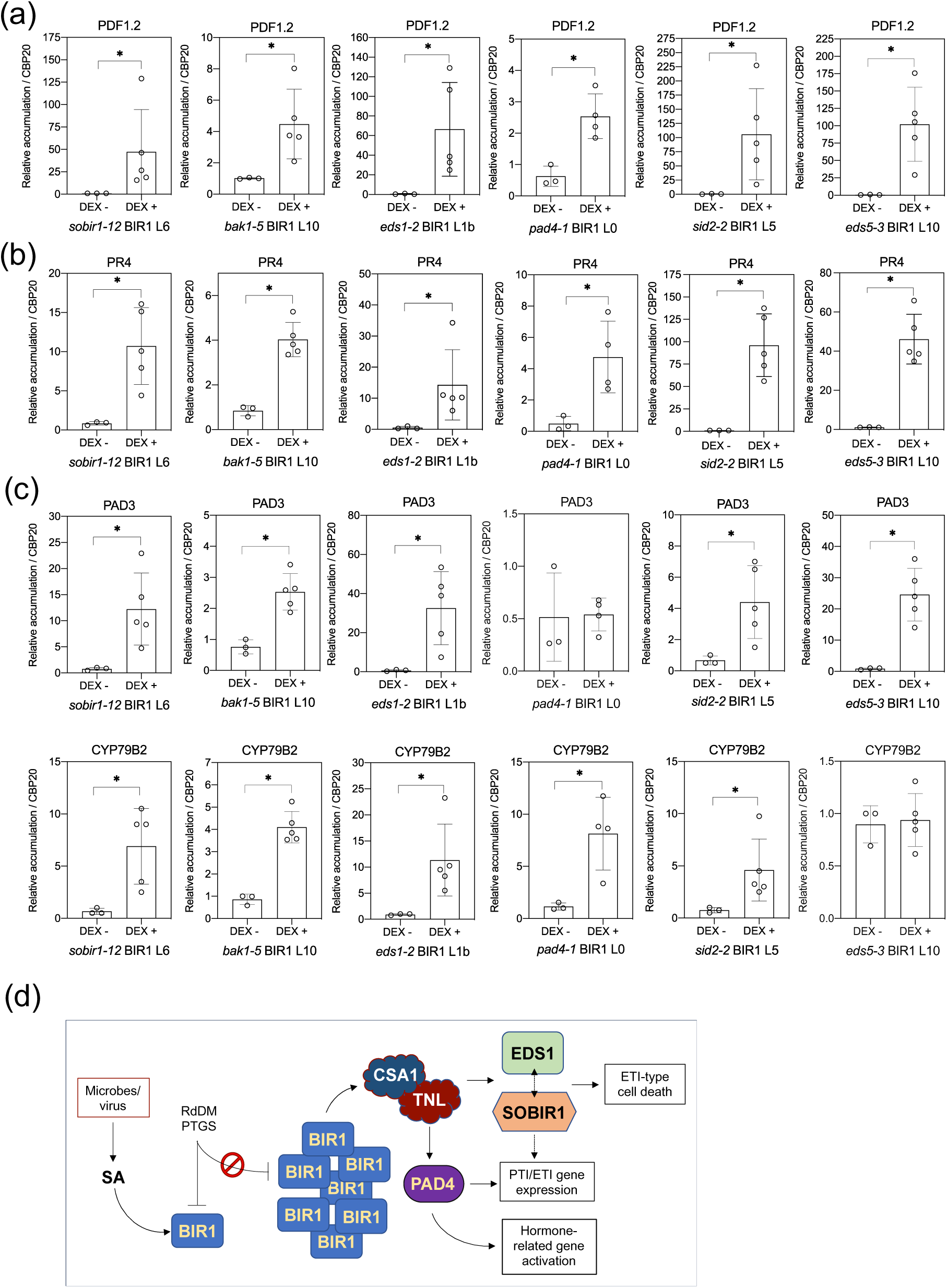
Expression analyses of phytohormone-related genes in *BIR1* overexpressing plants. **(a, b, c)** RT-qPCR analysis of *PDF1.2* (a), *PR4* (b), *PAD3* and *CYP79B2* (c) transcripts in *sobir1-12* BIR1 L6, *bak1-*5 BIR1 L10, *eds1-2* BIR1 L1b, *pad4-1* BIR1 L0, *sid2-2* BIR1 L5 and *eds5-3* BIR1 L10 mutant lines. Measures were done in mock-treated (DEX-) versus DEX-treated (DEX+) using the same set of replicates from each genotype. Values from biologically independent samples were normalized to the *CBP20* internal control, and related to mock-treated plants, set to a value of 1. Data are mean ± SD analyzed by Mann-Whitney test; *, *p* <0.05. **(d)** Tentative working model for BIR1 homeostasis. SA-mediated transcription of *BIR1* is regulated by RNA-directed DNA methylation (RdDM) and post-transcriptional RNA silencing (PTGS) (Guzman-Benito *et al*., 2019). Loss of *BIR1* regulation leads to constitutive *BIR1* overexpression. *BIR1* overexpression is sensed by one or several TIR1-containing nucleotide-binding domain leucine-rich repeat receptors (TNLs), of which *CSA1* is a plausible candidate, to promote *PAD4*-dependent up-regulation of immune-related and defense hormone genes. TNL activation also leads to effector triggered immunity (ETI)-type cell death that requires *EDS1* and *SOBIR1*.

## 4 DISCUSSION

Plants respond to microbes and viruses by enhancing the expression of multiple genes that function as regulators or executors of defense programs (Nishad *et al*., 2020). Among them, *BIR1* was shown to be transcriptionally up-regulated during bacterial and oomycete infections, and also during viral infections in a SA-dependent manner (Gao *et al*., 2009; Guzman-Benito *et al*., 2019). The observation that *bir1-1* mutants have increased resistance to the virulent oomycete *Hyaloperonospora parasitica* Noco2 and to the RNA virus *Tobacco rattle virus* (TRV) suggests an active role of *BIR1* in plant immunity (Gao *et al*., 2009; Guzman-Benito *et al*., 2019). Previously we found that overexpression of *BIR1* in Arabidopsis using two independent expression system, a viral vector or a transgenic inducible system, caused senescence and cell death-like phenotypes. Intriguingly, these phenotypes were similar to those observed under *BIR1* depletion in knockout plants, which suggest that defects in *BIR1* regulation compromise the correct functioning of this gene. In order to investigate the genetic components responsible for *BIR1* overexpressing phenotypes, we used a DEX-inducible system to express a mCherry-tagged *BIR1* transgene in different Arabidopsis mutants defective in key regulators of PTI and ETI signal transduction pathways. The resulting transgenic plants from different genotypes accumulated BIR1-mCherry protein at comparable levels upon DEX treatment, indicating similar induction efficiency in our experimental setting. Furthermore, *BIR1* is triggered in the absence of immune elicitors or effectors that could otherwise initiate undesirable localized and systemic defense responses in the plant.

We found that the cell death-type morphological phenotypes observed upon *BIR1* overexpression was accompanied by the inappropriate overexpression of genes encoding receptors and co-receptors with key functions in the perception of ligands/effectors and in the regulation of defense signal transduction. The up-regulation of genes encoding TNL-resistance proteins, but not CNLs, is an important feature linked to BIR1 overexpression. TNL-based immunity is mediated by two distinct signaling modules, *EDS1*/*SAG101*/*NRG1* and *EDS1*/*PAD4*/*ADR1* (Sun *et al*., 2021). Here, we provide genetic evidence that the *BIR1* overexpression morphological phenotypes are partially dependent on *EDS1*, but not on *PAD4*. We propose that *EDS1*, perhaps in concert with *SAG101*, triggers cell death-like phenotypes upon TNL activation in *BIR1* overexpressing plants (Figure 8d). How TNL receptors are activated when *BIR1* is overexpressed is a question to resolve. A current model postulates that loss of *BIR1* function leads to the activation of hypothetical guarding R proteins, which further feed *PAD4*- and *SOBIR1*-dependent resistance pathways (Gao *et al*., 2009). Our results are coherent with BIR1 or a yet unknown BIR1/SERK complex being guarded by one or several *EDS1*-dependent TNL proteins. Therefore, overexpression of *BIR1* may promote TNL-mediated resistance pathways that require *EDS1* when no effectors or elicitors are present.

In addition to *EDS1*, we demonstrated that *SOBIR1* is necessary for *BIR1* overexpression phenotypes. The role of *SOBIR1* in the *BIR1* pathway is unclear. Although extensive investigation is required, S*OBIR1* may function as a specific co-regulator of the *EDS1*-dependent responses triggered by *BIR1* overexpression, as part of the same signaling pathway (Figure 8d). This idea is inspired by a recent work showing that the cell-surface RLP23 receptor uses SOBIR1 to form a supramolecular complex with a EDS1/PAD4/ADR1 node in a ligand-independent manner to mediate defense signaling cascades that are common to both surface PRR (PTI) and NLR (ETI) receptors (Pruitt *et al*., 2021b).

The TNL protein CSA1 senses perturbances of BIR3, BAK1 or BIR3/BAK1 complexes leading to ETI-type autoimmune responses (Schulze *et al*., 2022). *CSA1* did not show responsiveness to *BIR1* overexpression in our assay, but it has been shown to interact with BIR1 (Schulze *et al*., 2022). It will be interesting to see if CSA1 also guards BIR1 integrity, perhaps in cooperation with SOBIR1 and/or other SERKs (Figure 8d). Finally, phenotypes linked to *BIR1* overexpression were apparently unaffected by the *bak1-5* mutation, which suggests that other SERK family members can compensate for the loss of *BAK1* in PTI/ETI signaling.

We observed that the upregulation of *FRK1*, *WRKY29* and *RBOHD* due to *BIR1* overexpression was compromised by mutations in *SOBIR1* and *PAD4,* but it was independent of *EDS1*. Since *EDS1*, *PAD4* and *SOBIR1* are important for both cell surface PRR receptor- and TNL-mediated signaling and resistance (Pruitt *et al*., 2021b; Tian *et al*., 2021), we will need additional experimentation to establish if changes in PTI gene expression involve PTI-dependent pathways, TNL activation or other unknown mechanisms yet to be determined. Our study also reveals significant alterations in plant hormone gene expression caused by *BIR1* overexpression. Interestingly, genes related to the plant hormones ET, JA, CA, and SA, which play a central role in regulating immune responses, remained up-regulated in *eds1-2* BIR1 and *sobir1-12* BIR1 backgrounds, in which phenotypes were partially rescued. This suggests that i) regulation of hormone homeostasis does not require *SOBIR1*- and/or *EDS1*-mediated signaling pathways, and that ii) the hormonal imbalance in BIR1 overexpressing plants has little, if any, influence on the phenotypes.

The Arabidopsis heterotrimeric G protein AGB1 (SOBIR2) is a common signaling component of PTI mediated by different RLK-PRRs, and functions in parallel with PAD4 downstream of SOBIR1 to regulate cell death and pathogen resistance in *bir1-1* Arabidopsis mutants (Liu *et al*., 2013; Liang *et al*., 2016). Similarly, Gγ-proteins AGG1/AGG2 have been described as positive regulators of cell death and constitutive defense phenotypes in *bir1-1* mutants (Liu *et al*., 2013). *AGB1* and *AGG2* were up-regulated in *BIR1* overexpressing plants. These findings bring up the idea that AGB1 and AGG1/2 could also function downstream of BIR1 to regulate *EDS1*-mediated cell death when BIR1 integrity is disrupted. Equally important, the *BIR1*-responsive GPA1 and XLG, in association with Gβγ modules, also have crucial roles in multiple growth and developmental processes (Wang *et al*., 2023). Thus, it is conceivable that overactivation of these heterotrimeric G proteins due to the overexpression of *BIR1* may cause indirect effects on plant development. Further studies are needed to test these plausible hypotheses and their applicability to other plant species. In addition, identification of *BIR1* interactors shall increase our understanding of the underlying mechanisms and components of the *BIR1*-mediated regulation of immunity.

An interesting topic in this work is whether DEX treatment may be synergistically responsible for the morphological phenotypes observed upon *BIR1* overexpression in the inducible system. However, we find this possibility unconvincing. Although DEX may cause moderate plant growth retardation in an age-dependent manner in our system, cell death/senescence phenotypes were observed in none of the DEX-treated WT and mutant plants (without the BIR1-mCherry cassette) or in transgenic plants expressing a mCherry-coding construct. In addition, the cell death phenotypes described using the DEX-inducible system were similar to those reported in WT Arabidopsis plants infected with a viral vector that overexpressed *BIR1* fused to a HA protein tag. In these plants, phenotypes were normally stronger in infected tissue accumulating high levels of HA-tagged BIR1 (Guzman-Benito *et al*., 2019). Thus, it is unlikely that the severe phenotypes observed in BIR1 L9 plants were an artifact caused by either DEX treatment or mCherry overexpression, or by a synergistic interaction between DEX, *BIR1* overexpression and tagged proteins (Guzman-Benito *et al*., 2019). Because overexpression of BIR1-mCherry has the same phenotypical outputs as HA-tagged BIR1 (Guzman-Benito *et al*., 2019), and HA-tagging of *BIR1* produces a functional protein that is able to fully complement *bir1-1* autoimmunity in Arabidopsis (Gao *et al*., 2009), we suspect that the mCherry-tagging does not compromise *BIR1* function in our transgenic setup. In summary, even though some adverse effects due to long DEX treatments cannot be fully discarded in our experimental system, specific effects linked to *BIR1* overexpression could be distinguished from other potential combinatorial effects resulting from expressing an mCherry-tagged version of *BIR1* in an activated glucocorticoid-inducible background (Kang *et al*., 1999).

In conclusion, we investigated the implications of *BIR1* homeostasis on the immune response in Arabidopsis. Overexpression of *BIR1* results in cell death phenotypes that resemble the previously reported in loss-of-function *bir1-1* mutants. Furthermore, *BIR1* overexpression led to massive transcriptomic changes affecting multiple defense-related genes. We show that both *EDS1* and *SOBIR1* are necessary for the cell death phenotypes observed in *BIR1* overexpressing plants, while *PAD4* and *BAK1* are not. We hypothesize that BIR1, previously thought to represent a negative regulator of plant immunity, is a guarded protein surveilled by unknown TNLs, with SOBIR1 working alongside EDS1 to transduce signals downstream of the TNLs (Figure 8d).

## ACKNOWLEDGEMENTS

We thank members of the César Llave lab and to my colleagues Manfred Heinlein (Institut de Biologie Moléculaire des Plantes, CNRS, Strasbourg, France), Rosa Lozano-Durán, Birgit Kemmerling, Thorsten Nürnberger (Center of Plant Molecular Biology, University of Tübingen, Tübingen, Germany) and Francisco Tenllado (Centro de Investigaciones Biológicas Margarita Salas, CSIC, Madrid, Spain) for helpful discussions. We thank Monica Fontenla for technical support.

## AUTHOR CONTRIBUTIONS

CL and IGB conceived and designed the study. CL, IGB, CR and IP performed expression assays and transgenics. CL and IGB analyzed the data. CL wrote the article.

## Data availability

The data supporting this study’s findings are available from the corresponding author upon reasonable request.

## SUPPORTING INFORMATION

### SUPPORTING FIGURES

**Figure S1.**
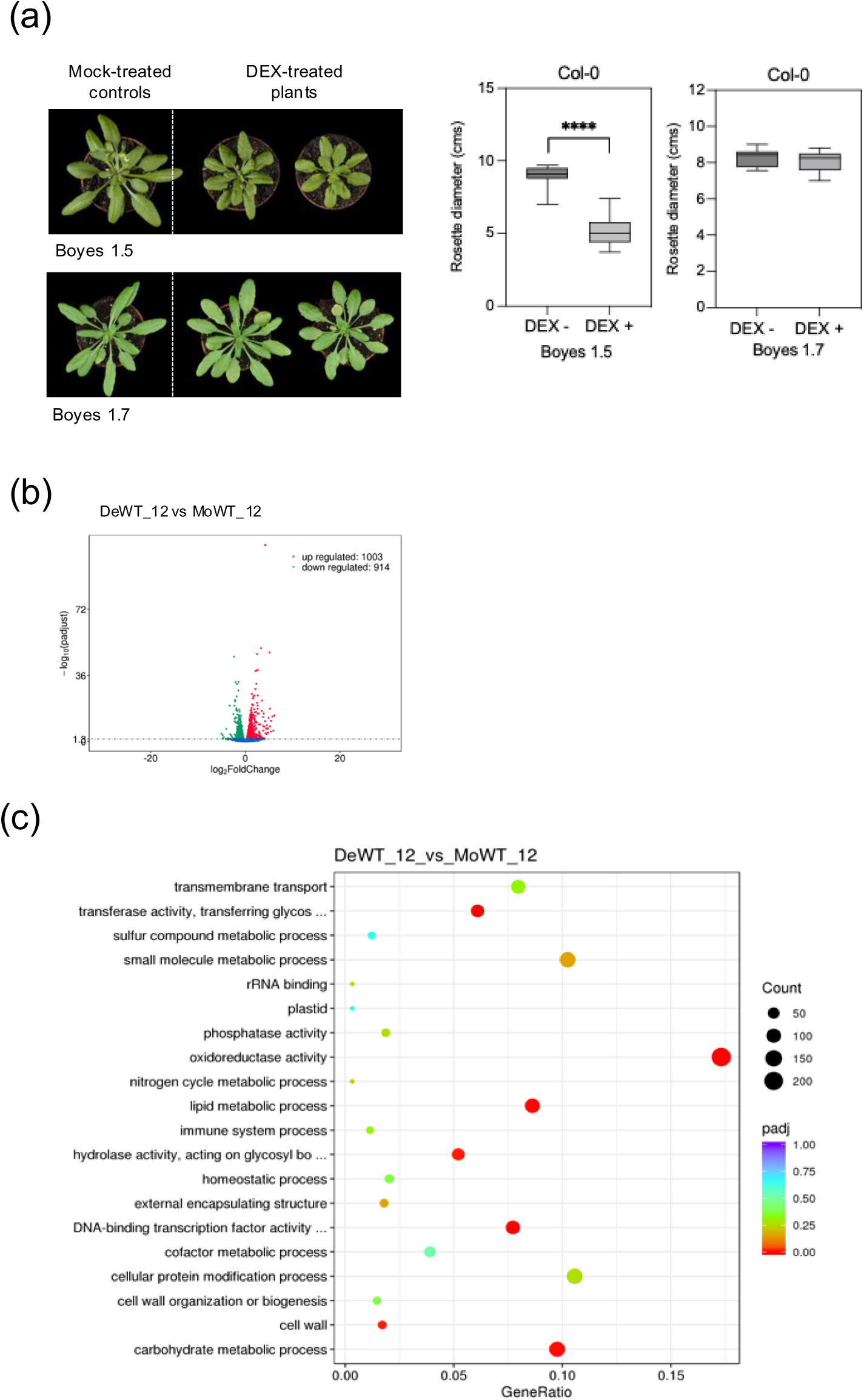
Transcriptomic profiling of dexamethasone (DEX)-treated wild-type (WT) Col-0 plants. **(a)** Morphology of representative WT Arabidopsis plants after 10-12 days of 30 µM DEX treatment. Mock-treated plants are shown as controls. DEX treatments started at the Boyes growth stages 1.5 or 1.7, as indicated. Rosette size (diameter) in WT (Boyes 1.5 and 1.7) after 12 days of DEX treatment relative to mock-treated controls is shown. Box and whisker plots showing 5-95 percentile were analyzed by Mann-Whitney test; ****, p <0.0001. **(b)** Volcano plots showing the distribution of differentially expressed genes (DEGs) (adjusted *p*-value ≤ 0.05). The x-axis shows the gene fold change, and y-axis plots the statistical significance (*p*-value) of the differences. Significantly up- and down-regulated genes are highlighted in red and green, respectively. Genes that are not differentially regulated are in blue. **(c)** Scatterplot of selected gene ontology (GO) terms of DEGs. GO terms between DEX-treated and mock-treated WT plants at 12 days of treatment (DeL9_ vs MoL9_) are shown. Dot size represents the count of different genes and the color indicates significance of the term enrichment. GO terms with adjusted *p*-value < 0.05 are significantly enriched. WT, wild-type Col-0; De, DEX-treated plants; Mo, Mock-treated plants; _12, 12 days of treatment.

**Figure S2.**
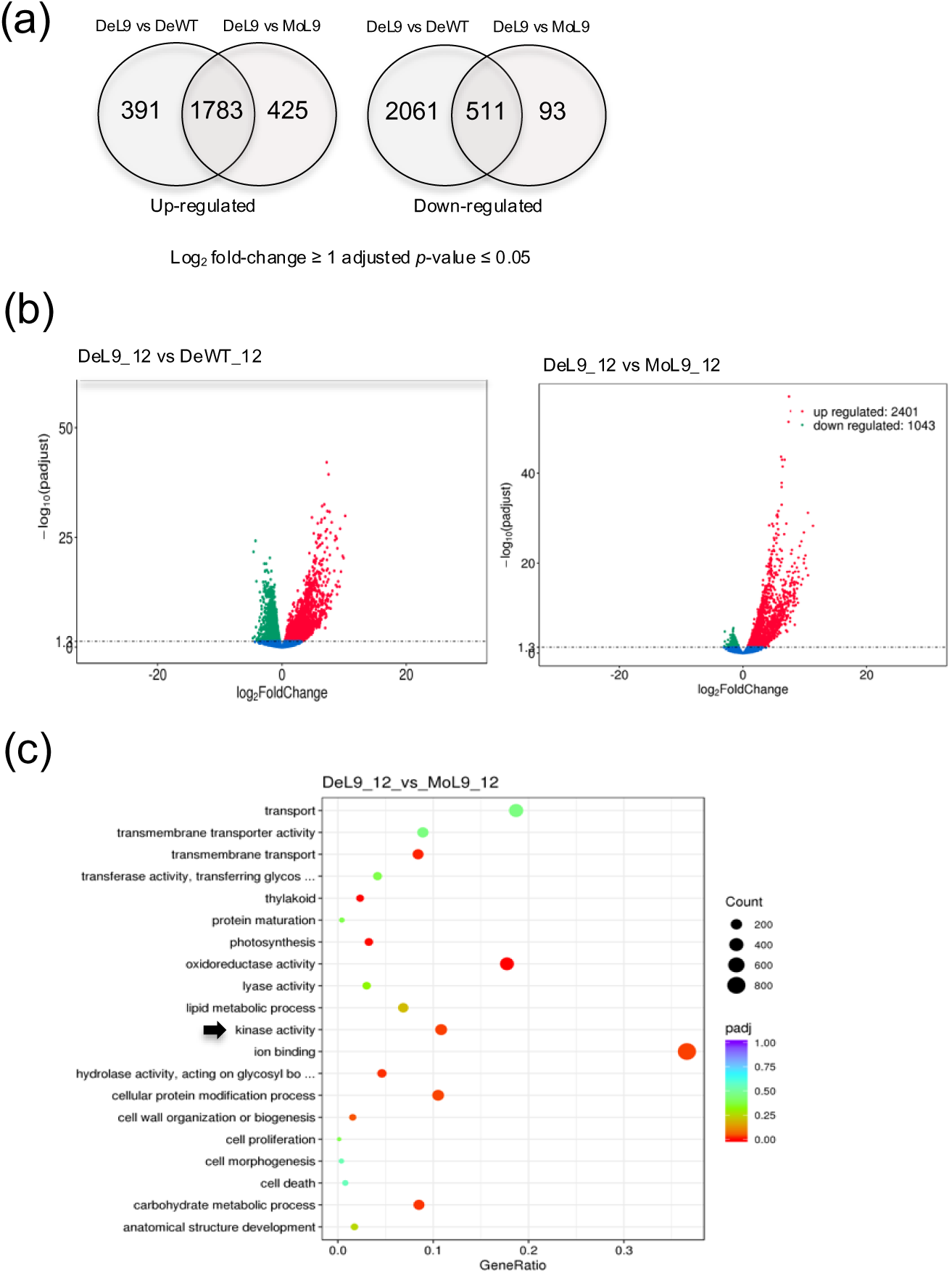
Transcriptomic profiling of *BIR1* expressing transgenic lines. **(a)** Number of up-or down-regulated transcripts, with a |log_2_ (Fold-Change)| ≥1 and adjusted *p*-value ≤0.05, in each pairwise comparison as indicated. **(b)** Volcano plots showing the distribution of DEGs (adjusted *p*-value ≤ 0.05). The x-axis shows the fold change in gene expression between samples, and y-axis shows the statistical significance of the differences. Significantly up- and down-regulated genes are highlighted in red and green, respectively. Genes that are not differentially regulated are in blue. **(c)** Scatterplot of selected gene ontology (GO) terms of DEGs. GO terms between wild-type (WT) and BIR1 L9 plants at 12 days after DEX-treatment (DeL9_ vs MoL9_) are shown. Dot size represents the count of different genes and the color indicates significance of the term enrichment. GO terms with adjusted *p*-value < 0.05 are significantly enriched. The kinase term (indicated by an arrow) was exclusively assigned to BIR1 L9 plants. WT, wild-type Col-0; L9, BIR1 L9 transgenic plants; De, DEX-treated plants; Mo, Mock-treated plants; _12, 12 days of treatment.

**Figure S3.**
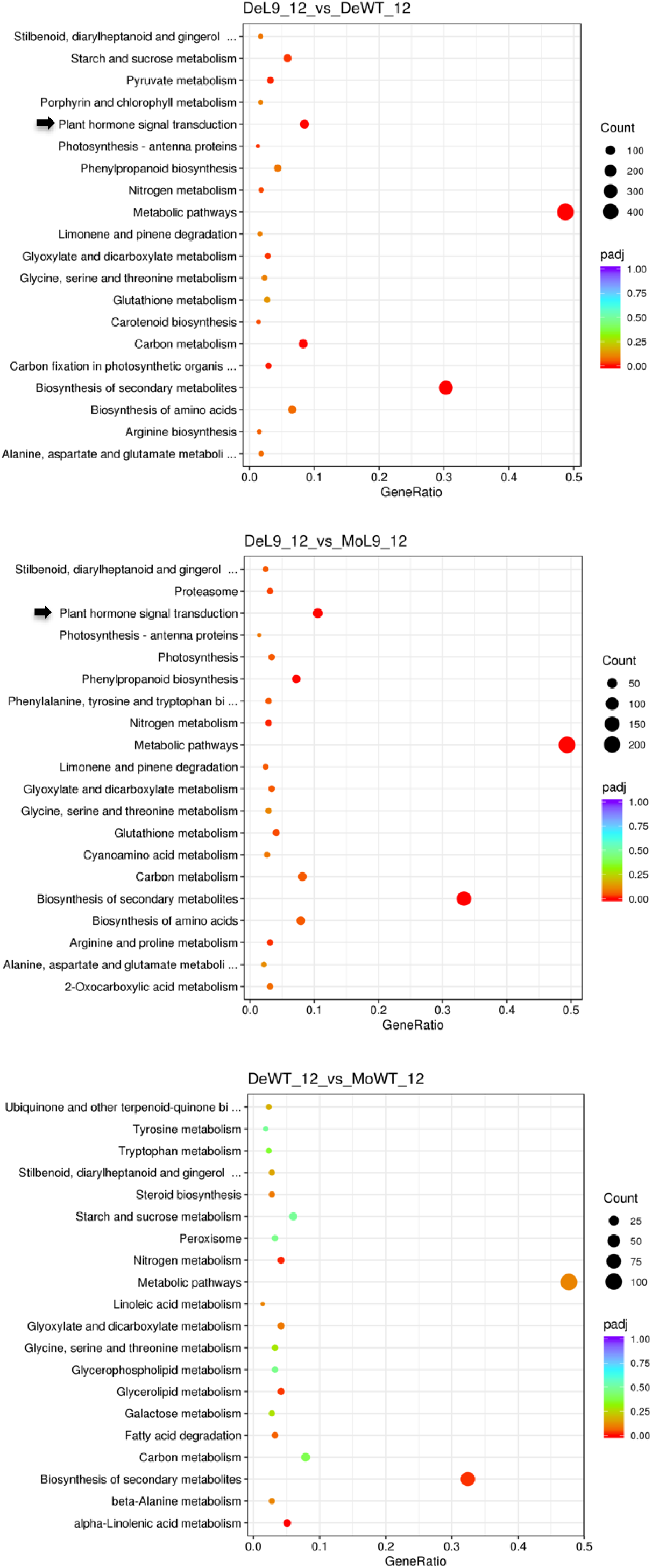
Enrichment analysis of Kyoto Encyclopedia of Genes and Genomes (KEGG) pathways using RNA-Seq data. Scatterplot of selected Kyoto Encyclopedia of Genes and Genomes (KEGG) pathways. Dot size represents the count of different genes and the color indicates significance of the term enrichment. Terms with adjusted *p*-value < 0.05 are significantly enriched. WT, wild-type Col-0; De, DEX-treated plants; Mo, Mock-treated plants; _12, 12 days of treatment.

**Figure S4.**
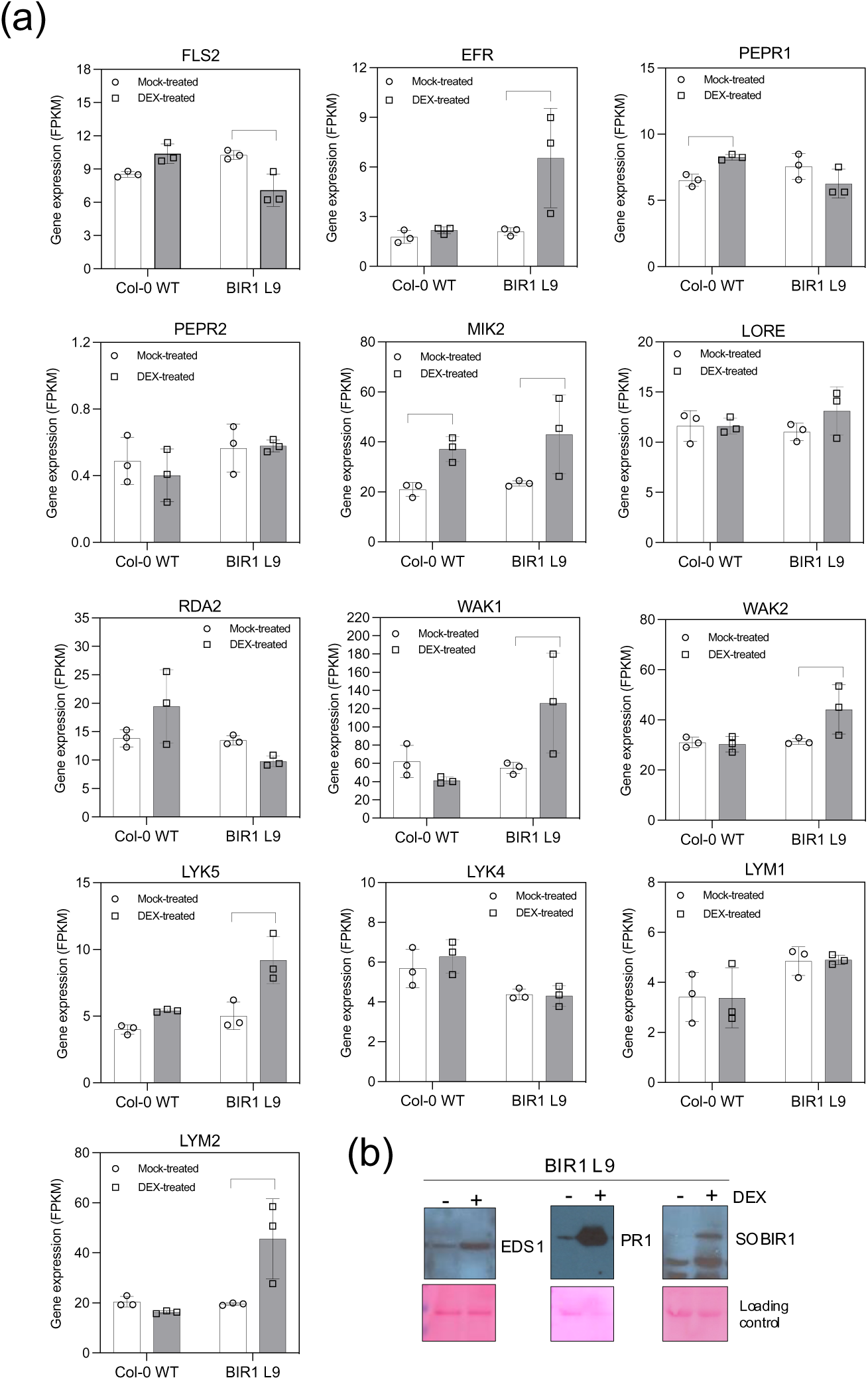
Differential expression of cell-surface pattern recognition receptors (PRR)-coding genes in *BIR1* overexpressing plants. **(a)** Relative expression levels are represented by the number of Fragments Per Kilobase of transcript sequence per Millions of base-pairs sequenced (FPKM) in each of the three independent samples per condition used for RNA-Seq. Significant differences between dexamethasone (DEX)- and mock-treated samples were analyzed using two-way ANOVA followed by Sidak’s multiple comparison test; *, adjusted *p*-value < 0.05. **(b)** Western blot analysis of EDS1, SOBIR1 and PR1 using specific antibodies in two independent sets of BIR1 L9 samples. Plants were mock-treated (-) or DEX-treated (+) for 12 days. Protein loading control by Ponceau S is shown.

**Figure S5.**
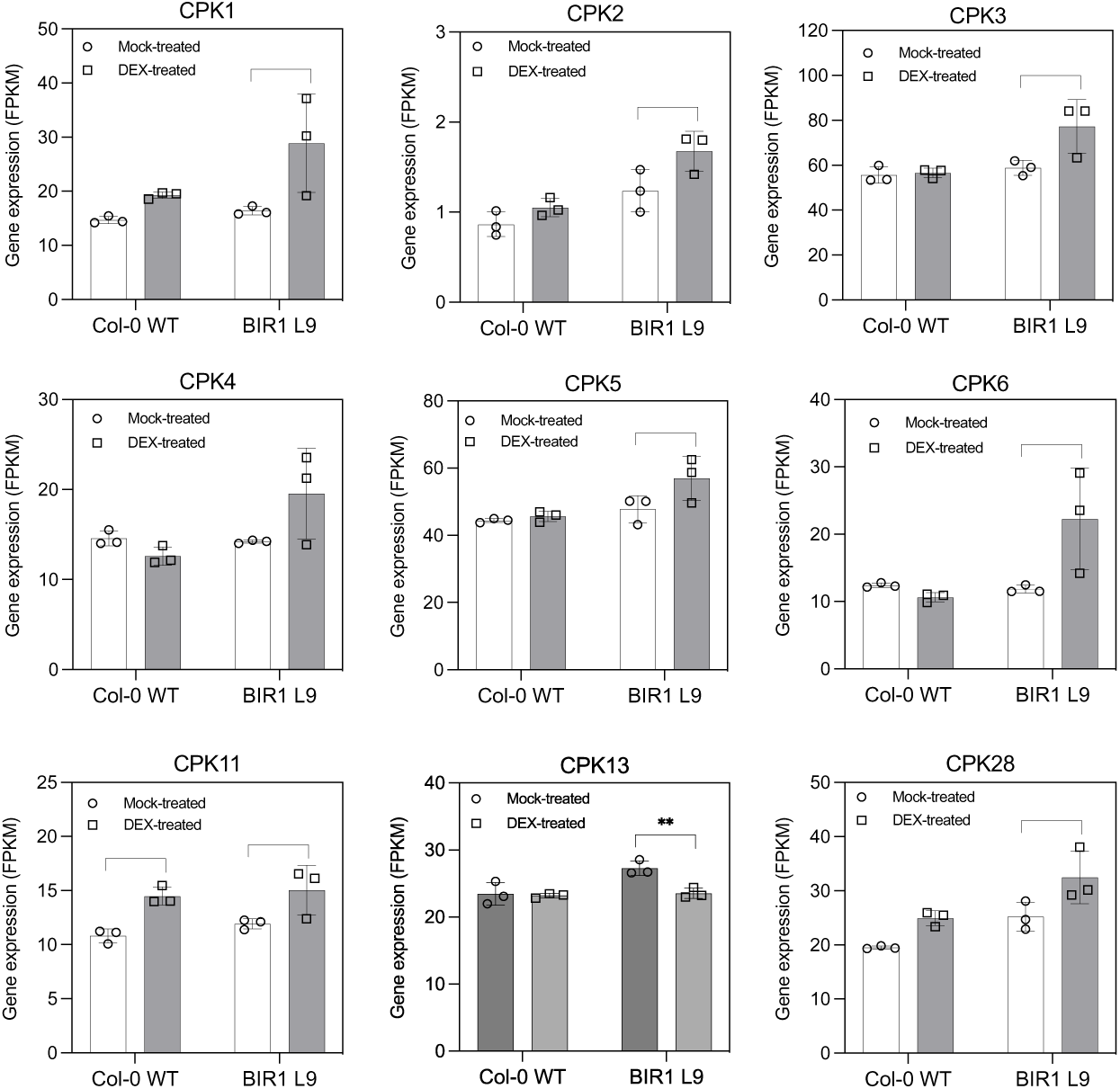
Differential expression of calcium-dependent protein kinases (CPK)-coding genes in *BIR1* overexpressing plants. Relative expression levels are represented by the number of Fragments Per Kilobase of transcript sequence per Millions of base-pairs sequenced (FPKM) in each of the three independent samples per condition used for RNA-Seq. Significant differences between dexamethasone (DEX)- and mock-treated samples were analyzed using two-way ANOVA followed by Sidak’s multiple comparison test; **, adjusted *p*-value < 0.01, *< 0.05.

**Figure S6.**
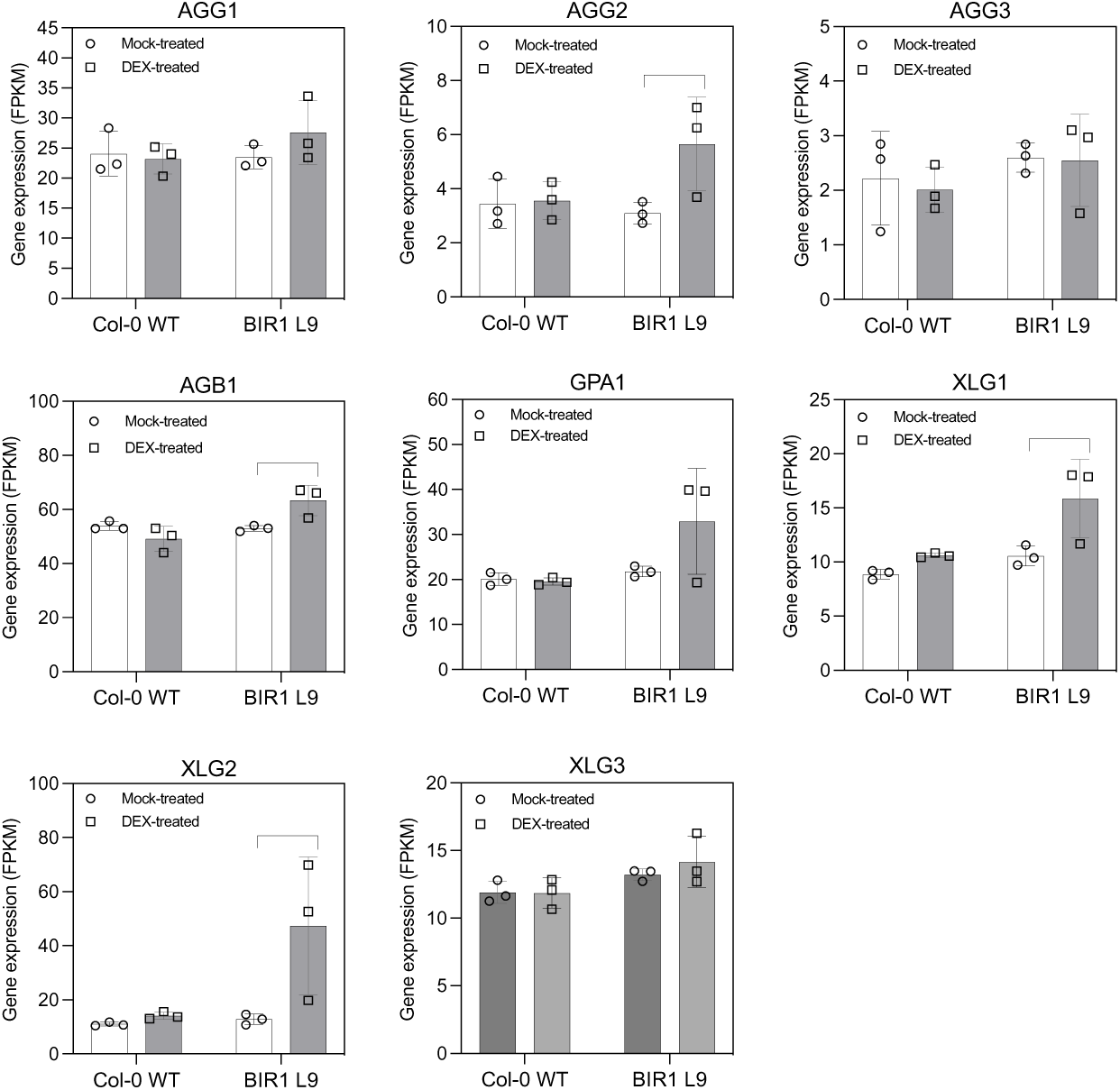
Differential expression of *HETEROTRIMERIC G PROTEIN* genes in *BIR1* overexpressing plants. Relative expression levels are represented by the number of Fragments Per Kilobase of transcript sequence per Millions of base-pairs sequenced (FPKM) in each of the three independent samples per condition used for RNA-Seq. Significant differences between dexamethasone (DEX)- and mock-treated samples were analyzed using two-way ANOVA followed by Sidak’s multiple comparison test; *, adjusted *p*-value < 0.05.

**Figure S7.**
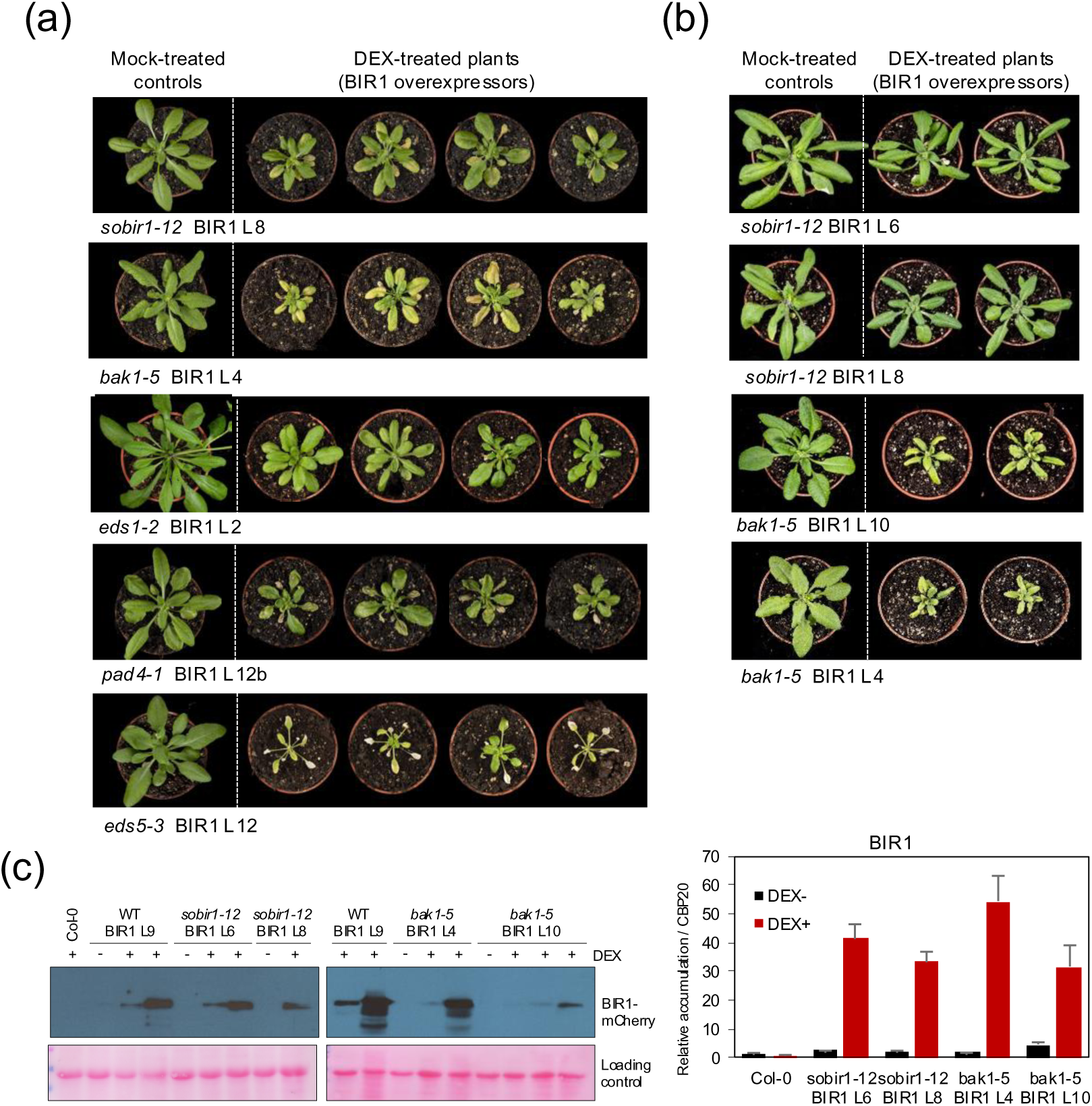
Morphological and growth defects in independent dexamethasone (DEX)-treated *BIR1* overexpressing plants of different immune mutant genetic backgrounds. **(a)** Morphology of representative DEX-inducible BIR1-mCherry transgenic Arabidopsis lines in *sobir1-12* (*sobir1-12* BIR1 L8), *bak1-*5 (*bak1-*5 BIR1 L4), *eds1-*2 (*eds1-*2 BIR1 L2), *pad4-1* (*pad4-1* BIR1 L12b) and *eds5-3* (*eds5-3* BIR1 L12) backgrounds. Photographs were taken after 10-12 days of 30 µM DEX treatment. Mock-treated plants are shown as controls. **(b)** Morphology and *BIR1* expression levels of DEX-treated and mock-treated *sobir1-12* BIR1 (L6, L8) and *bak1-5* BIR1 (L4, L10) plants as observed in a different repetition of the experiment. **(c)** Western blot analysis of BIR1-mCherry in the genotypes indicated. Plants were mock-treated (-) or DEX-treated (+) for 12 days. Protein loading control by Ponceau S is shown (left panel). qRT-PCR analysis showed comparable *BIR1* transcript levels in both genotypes. Each sample consisted of RNA pooled from six different plants (right panel).

**Figure S8.**
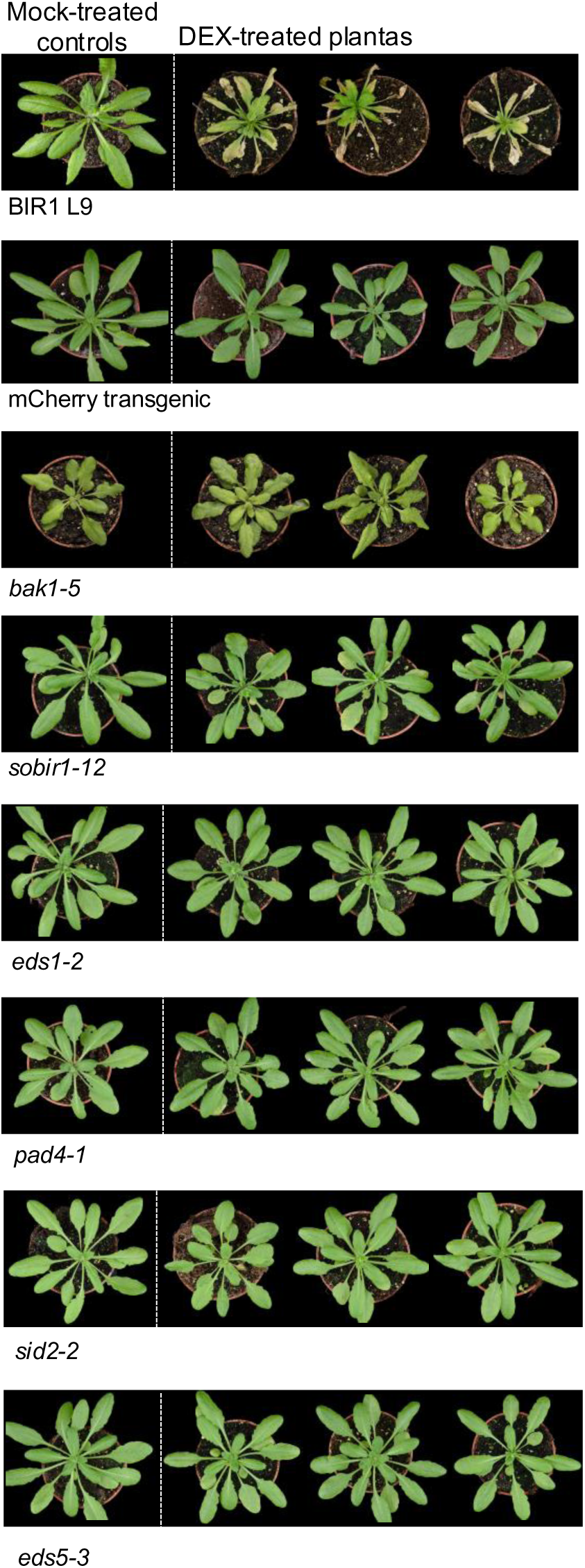
Morphological phenotypes of dexamethasone (DEX)-treated immune mutant genetic backgrounds and mCherry transgenic plants. Morphology of BIR1 L9, *sobir1-12*, *bak1-5, eds1-2*, *pad4-1*, *sid2-2* and *eds5-3* mutant plants and mCherry transgenic Arabidopsis after 12 days of treatment with 30 µM. Mock-treated plants of each genotype are shown as controls.

### SUPPORTING TABLES

**Table S1.** List of primers

**Table S2** List of expressed genes. Values represent the total read counts/ Fragments Per Kilobase of transcript sequence per Millions of base-pairs sequenced (FRKM) values of each biological replicate (a,b,c) per condition and their corresponding row mean values. De (DEX-treated plants), Mo (Mock-treated plants), WT (wild-type plants), L9 (BIR1 L9 transgenic plants), _4 (4 days of treatment), _12 (12 days of treatment)

**Table S3** List of differentially expressed genes (DEG. padjust value < 0.05). Values represent the mean Fragments Per Kilobase of transcript sequence per Millions of base-pairs sequenced (FPKM) values from three biological replicates per condition. De (DEX-treated plants), Mo (Mock-treated plants), WT (wild-type plants), L9 (BIR1 L9 transgenic plants), _12 (12 days of treatment).

**Table S4** List of differentially expressed genes (DEG. |Log_2_fold change > 1| padjust value < 0.01). Values represent the mean Fragments Per Kilobase of transcript sequence per Millions of base-pairs sequenced (FPKM) values from three biological replicates per condition. De (DEX-treated plants), Mo (Mock-treated plants), WT (wild-type plants), L9 (BIR1 L9 transgenic plants), _12 (12 days of treatment).

**Table S5** Enrichment analysis of Gene Ontology (GO) terms. CC (cellular component), BP (biological process) and MF (molecular function). Gene Ratio is the ratio of DEGs to all genes for this GO term. Bg Ratio is the ratio of all genes concerning this GO term to all genes in background GO database. Count is the number of differentially expressed genes concerning this GO term. Adjusted p-value < 0.05 means significant enrichment. De (DEX-treated plants), Mo (Mock-treated plants), WT (wild-type plants), L9 (BIR1 L9 transgenic plants), _12 (12 days of treatment).

**Table S6** Kyoto Encyclopedia of Genes and Genomes (KEGG) pathway enrichment analysis. Gene Ratio is the ratio of DEGs to all genes for this pathway. Bg Ratio is the ratio of all genes concerning this KEGG term to all genes in background KEGG database. Count is the number of differentially expressed genes concerning this term. Adjusted p-value < 0.05 means significant enrichment. De (DEX-treated plants), Mo (Mock-treated plants), WT (wild-type plants), L9 (BIR1 L9 transgenic plants), _12 (12 days of treatment).

**Table S7** List of differentially expressed genes (DEG. padjust value < 0.05) within functional categories. Values represent the mean Fragments Per Kilobase of transcript sequence per Millions of base-pairs sequenced (FPKM) values from three biological replicates per condition. De (DEX-treated plants), Mo (Mock-treated plants), WT (wild-type plants), L9 (BIR1 L9 transgenic plants), _12 (12 days of treatment).

**Table S8** Statistical analysis of enrichment of functional gene groups in plants overexpressing BIR1 after 12 days of DEX treatment relative to non-expressing controls using Two-tailed Fisher’s exact tests. De (DEX-treated plants), Mo (Mock-treated plants), WT (wild-type plants), L9 (BIR1 L9 transgenic plants). LRR, leucine-rich repeat; RLK, receptor-like kinase; CRK, cysteine-rich RLKs; RLP, receptor-like protein; TNL, TIR1-nucleotide-binding domain LRR immune receptors.

## REFERENCES

Albert I, Zhang L, Bemm H, Nurnberger T. 2019. Structure-Function Analysis of Immune Receptor AtRLP23 with Its Ligand nlp20 and Coreceptors AtSOBIR1 and AtBAK1. Mol Plant Microbe Interact 32(8): 1038–1046.

Belkhadir Y, Jaillais Y, Epple P, Balsemao-Pires E, Dangl JL, Chory J. 2012. Brassinosteroids modulate the efficiency of plant immune responses to microbe-associated molecular patterns. Proc Natl Acad Sci U S A 109(1): 297–302.

Bi G, Zhou Z, Wang W, Li L, Rao S, Wu Y, Zhang X, Menke FLH, Chen S, Zhou JM. 2018. Receptor-Like Cytoplasmic Kinases Directly Link Diverse Pattern Recognition Receptors to the Activation of Mitogen-Activated Protein Kinase Cascades in Arabidopsis. Plant Cell 30(7): 1543–1561.

Boudsocq M, Willmann MR, McCormack M, Lee H, Shan L, He P, Bush J, Cheng SH, Sheen J. 2010. Differential innate immune signalling via Ca(2+) sensor protein kinases. Nature 464(7287): 418–422.

Boutrot F, Zipfel C. 2017. Function, Discovery, and Exploitation of Plant Pattern Recognition Receptors for Broad-Spectrum Disease Resistance. Annu Rev Phytopathol 55: 257–286.

Boyes DC, Zayed AM, Ascenzi R, McCaskill AJ, Hoffman NE, Davis KR, Gorlach J. 2001. Growth stage-based phenotypic analysis of Arabidopsis: a model for high throughput functional genomics in plants. Plant Cell 13(7): 1499–1510.

Brodersen P, Petersen M, Bjorn Nielsen H, Zhu S, Newman MA, Shokat KM, Rietz S, Parker J, Mundy J. 2006. Arabidopsis MAP kinase 4 regulates salicylic acid- and jasmonic acid/ethylene-dependent responses via EDS1 and PAD4. Plant J 47(4): 532–546.

Brutus A, Sicilia F, Macone A, Cervone F, De Lorenzo G. 2010. A domain swap approach reveals a role of the plant wall-associated kinase 1 (WAK1) as a receptor of oligogalacturonides. Proc Natl Acad Sci U S A 107(20): 9452–9457.

Cao Y, Liang Y, Tanaka K, Nguyen CT, Jedrzejczak RP, Joachimiak A, Stacey G. 2014. The kinase LYK5 is a major chitin receptor in Arabidopsis and forms a chitin-induced complex with related kinase CERK1. Elife 3.

Castro B, Citterico M, Kimura S, Stevens DM, Wrzaczek M, Coaker G. 2021. Stress-induced reactive oxygen species compartmentalization, perception and signalling. Nat Plants 7(4): 403–412.

Chen Z. 2001. A superfamily of proteins with novel cysteine-rich repeats. Plant Physiol 126(2): 473–476.

Clough SJ, Bent AF. 1998. Floral dip: a simplified method for Agrobacterium-mediated transformation of Arabidopsis thaliana. Plant J 16(6): 735–743.

Couto D, Zipfel C. 2016. Regulation of pattern recognition receptor signalling in plants. Nat Rev Immunol 16(9): 537–552.

Cui H, Gobbato E, Kracher B, Qiu J, Bautor J, Parker JE. 2017. A core function of EDS1 with PAD4 is to protect the salicylic acid defense sector in Arabidopsis immunity. New Phytol 213(4): 1802–1817.

Dominguez-Ferreras A, Kiss-Papp M, Jehle AK, Felix G, Chinchilla D. 2015. An Overdose of the Arabidopsis Coreceptor BRASSINOSTEROID INSENSITIVE1-ASSOCIATED RECEPTOR KINASE1 or Its Ectodomain Causes Autoimmunity in a SUPPRESSOR OF BIR1-1-Dependent Manner. Plant Physiol 168(3): 1106–1121.

Faulkner C, Petutschnig E, Benitez-Alfonso Y, Beck M, Robatzek S, Lipka V, Maule AJ. 2013. LYM2-dependent chitin perception limits molecular flux via plasmodesmata. Proc Natl Acad Sci U S A 110(22): 9166–9170.

Fernandez-Calvino L, Guzman-Benito I, Del Toro FJ, Donaire L, Castro-Sanz AB, Ruiz-Ferrer V, Llave C. 2016. Activation of senescence-associated Dark-inducible (DIN) genes during infection contributes to enhanced susceptibility to plant viruses. Mol Plant Pathol 17(1): 3–15.

Freh M, Gao JL, Petersen M, Panstruga R. 2022. Plant autoimmunity-fresh insights into an old phenomenon. Plant Physiology 188(3): 1419–1434.

Gao M, Wang X, Wang D, Xu F, Ding X, Zhang Z, Bi D, Cheng YT, Chen S, Li X, et al. 2009. Regulation of cell death and innate immunity by two receptor-like kinases in Arabidopsis. Cell Host Microbe 6(1): 34–44.

Gust AA, Pruitt R, Nurnberger T. 2017. Sensing Danger: Key to Activating Plant Immunity. Trends Plant Sci 22(9): 779–791.

Guzman-Benito I, Donaire L, Amorim-Silva V, Vallarino JG, Esteban A, Wierzbicki AT, Ruiz-Ferrer V, Llave C. 2019. The immune repressor BIR1 contributes to antiviral defense and undergoes transcriptional and post-transcriptional regulation during viral infections. New Phytol 224(1): 421–438.

Halter T, Imkampe J, Mazzotta S, Wierzba M, Postel S, Bucherl C, Kiefer C, Stahl M, Chinchilla D, Wang X, et al. 2014. The leucine-rich repeat receptor kinase BIR2 is a negative regulator of BAK1 in plant immunity. Curr Biol 24(2): 134–143.

He ZH, Fujiki M, Kohorn BD. 1996. A cell wall-associated, receptor-like protein kinase. J Biol Chem 271(33): 19789–19793.

Hohmann U, Lau K, Hothorn M. 2017. The Structural Basis of Ligand Perception and Signal Activation by Receptor Kinases. Annu Rev Plant Biol 68: 109–137.

Hou S, Liu D, Huang S, Luo D, Liu Z, Xiang Q, Wang P, Mu R, Han Z, Chen S, et al. 2021. The Arabidopsis MIK2 receptor elicits immunity by sensing a conserved signature from phytocytokines and microbes. Nat Commun 12(1): 5494.

Huang W, Wang Y, Li X, Zhang Y. 2020. Biosynthesis and Regulation of Salicylic Acid and N-Hydroxypipecolic Acid in Plant Immunity. Mol Plant 13(1): 31–41.

Imkampe J, Halter T, Huang S, Schulze S, Mazzotta S, Schmidt N, Manstretta R, Postel S, Wierzba M, Yang Y, et al. 2017. The Arabidopsis Leucine-Rich Repeat Receptor Kinase BIR3 Negatively Regulates BAK1 Receptor Complex Formation and Stabilizes BAK1. Plant Cell 29(9): 2285–2303.

Jirage D, Tootle TL, Reuber TL, Frost LN, Feys BJ, Parker JE, Ausubel FM, Glazebrook J. 1999. Arabidopsis thaliana PAD4 encodes a lipase-like gene that is important for salicylic acid signaling. Proc Natl Acad Sci U S A 96(23): 13583–13588.

Jubic LM, Saile S, Furzer OJ, El Kasmi F, Dangl JL. 2019. Help wanted: helper NLRs and plant immune responses. Curr Opin Plant Biol 50: 82–94.

Kadota Y, Sklenar J, Derbyshire P, Stransfeld L, Asai S, Ntoukakis V, Jones JD, Shirasu K, Menke F, Jones A, et al. 2014. Direct regulation of the NADPH oxidase RBOHD by the PRR-associated kinase BIK1 during plant immunity. Mol Cell 54(1): 43–55.

Kang HG, Fang YW, Singh KB. 1999. A glucocorticoid-inducible transcription system causes severe growth defects in and induces defense-related genes. Plant Journal 20(1): 127–133.

Kato H, Nemoto K, Shimizu M, Abe A, Asai S, Ishihama N, Matsuoka S, Daimon T, Ojika M, Kawakita K, et al. 2022. Recognition of pathogen-derived sphingolipids in Arabidopsis. Science 376(6595): 857–860.

Korner CJ, Klauser D, Niehl A, Dominguez-Ferreras A, Chinchilla D, Boller T, Heinlein M, Hann DR. 2013. The immunity regulator BAK1 contributes to resistance against diverse RNA viruses. Mol Plant Microbe Interact 26(11): 1271–1280.

Krol E, Mentzel T, Chinchilla D, Boller T, Felix G, Kemmerling B, Postel S, Arents M, Jeworutzki E, Al-Rasheid KA, et al. 2010. Perception of the Arabidopsis danger signal peptide 1 involves the pattern recognition receptor AtPEPR1 and its close homologue AtPEPR2. J Biol Chem 285(18): 13471–13479.

Li L, Li M, Yu L, Zhou Z, Liang X, Liu Z, Cai G, Gao L, Zhang X, Wang Y, et al. 2014. The FLS2-associated kinase BIK1 directly phosphorylates the NADPH oxidase RbohD to control plant immunity. Cell Host Microbe 15(3): 329–338.

Li P, Lu YJ, Chen H, Day B. 2020. The Lifecycle of the Plant Immune System. CRC Crit Rev Plant Sci 39(1): 72–100.

Liang X, Ding P, Lian K, Wang J, Ma M, Li L, Li L, Li M, Zhang X, Chen S, et al. 2016. Arabidopsis heterotrimeric G proteins regulate immunity by directly coupling to the FLS2 receptor. Elife 5: e13568.

Liang X, Zhou JM. 2018. Receptor-Like Cytoplasmic Kinases: Central Players in Plant Receptor Kinase-Mediated Signaling. Annu Rev Plant Biol 69: 267–299.

Liu J, Ding P, Sun T, Nitta Y, Dong O, Huang X, Yang W, Li X, Botella JR, Zhang Y. 2013. Heterotrimeric G proteins serve as a converging point in plant defense signaling activated by multiple receptor-like kinases. Plant Physiol 161(4): 2146–2158.

Liu Y, Huang X, Li M, He P, Zhang Y. 2016. Loss-of-function of Arabidopsis receptor-like kinase BIR1 activates cell death and defense responses mediated by BAK1 and SOBIR1. New Phytol 212(3): 637–645.

Luo X, Wu W, Liang Y, Xu N, Wang Z, Zou H, Liu J. 2020. Tyrosine phosphorylation of the lectin receptor-like kinase LORE regulates plant immunity. Embo J 39(4): e102856.

Lv Y, Yang N, Wu J, Liu Z, Pan L, Lv S, Wang G. 2016. New insights into receptor-like protein functions in Arabidopsis. Plant Signal Behav 11(7): e1197469.

McNellis TW, Mudgett MB, Li K, Aoyama T, Horvath D, Chua NH, Staskawicz BJ. 1998. Glucocorticoid-inducible expression of a bacterial avirulence gene in transgenic Arabidopsis induces hypersensitive cell death. Plant J 14(2): 247–257.

Mithoe SC, Menke FL. 2018. Regulation of pattern recognition receptor signalling by phosphorylation and ubiquitination. Curr Opin Plant Biol 45(Pt A): 162–170.

Nawrath C, Heck S, Parinthawong N, Metraux JP. 2002. EDS5, an essential component of salicylic acid-dependent signaling for disease resistance in Arabidopsis, is a member of the MATE transporter family. Plant Cell 14(1): 275–286.

Ngou BPM, Ahn HK, Ding P, Jones JDG. 2021. Mutual potentiation of plant immunity by cell-surface and intracellular receptors. Nature 592(7852): 110–115.

Nishad R, Ahmed T, Rahman VJ, Kareem A. 2020. Modulation of Plant Defense System in Response to Microbial Interactions. Front Microbiol 11: 1298.

Ouwerkerk PB, de Kam RJ, Hoge JH, Meijer AH. 2001. Glucocorticoid-inducible gene expression in rice. Planta 213(3): 370–378.

Parker JE, Holub EB, Frost LN, Falk A, Gunn ND, Daniels MJ. 1996. Characterization of eds1, a mutation in Arabidopsis suppressing resistance to Peronospora parasitica specified by several different RPP genes. Plant Cell 8(11): 2033–2046.

Pruitt RN, Gust AA, Nurnberger T. 2021a. Plant immunity unified. Nat Plants 7(4): 382–383.

Pruitt RN, Locci F, Wanke F, Zhang L, Saile SC, Joe A, Karelina D, Hua C, Frohlich K, Wan WL, et al. 2021b. The EDS1-PAD4-ADR1 node mediates Arabidopsis pattern-triggered immunity. Nature 598(7881): 495–499.

Quezada EH, Garcia GX, Arthikala MK, Melappa G, Lara M, Nanjareddy K. 2019. Cysteine-Rich Receptor-Like Kinase Gene Family Identification in the Phaseolus Genome and Comparative Analysis of Their Expression Profiles Specific to Mycorrhizal and Rhizobial Symbiosis. Genes (Basel) 10(1).

Rao S, Zhou Z, Miao P, Bi G, Hu M, Wu Y, Feng F, Zhang X, Zhou JM. 2018. Roles of Receptor-Like Cytoplasmic Kinase VII Members in Pattern-Triggered Immune Signaling. Plant Physiol 177(4): 1679–1690.

Rodriguez E, El Ghoul H, Mundy J, Petersen M. 2016. Making sense of plant autoimmunity and ‘negative regulators’. FEBS J 283(8): 1385–1391.

Schulze S, Yu L, Hua C, Zhang L, Kolb D, Weber H, Ehinger A, Saile SC, Stahl M, Franz-Wachtel M, et al. 2022. The Arabidopsis TIR-NBS-LRR protein CSA1 guards BAK1-BIR3 homeostasis and mediates convergence of pattern- and effector-induced immune responses. Cell Host Microbe 30(12): 1717–1731 e1716.

Schwessinger B, Roux M, Kadota Y, Ntoukakis V, Sklenar J, Jones A, Zipfel C. 2011. Phosphorylation-dependent differential regulation of plant growth, cell death, and innate immunity by the regulatory receptor-like kinase BAK1. PLoS Genet 7(4): e1002046.

Smakowska-Luzan E, Mott GA, Parys K, Stegmann M, Howton TC, Layeghifard M, Neuhold J, Lehner A, Kong J, Grunwald K, et al. 2018. An extracellular network of Arabidopsis leucine-rich repeat receptor kinases. Nature 553(7688): 342–346.

Stroud EA, Jayaraman J, Templeton MD, Rikkerink EHA. 2022. Comparison of the pathway structures influencing the temporal response of salicylate and jasmonate defence hormones in Arabidopsis thaliana. Front Plant Sci 13: 952301.

Sun X, Lapin D, Feehan JM, Stolze SC, Kramer K, Dongus JA, Rzemieniewski J, Blanvillain-Baufume S, Harzen A, Bautor J, et al. 2021. Pathogen effector recognition-dependent association of NRG1 with EDS1 and SAG101 in TNL receptor immunity. Nat Commun 12(1): 3335.

Swiderski MR, Birker D, Jones JD. 2009. The TIR domain of TIR-NB-LRR resistance proteins is a signaling domain involved in cell death induction. Mol Plant Microbe Interact 22(2): 157–165.

Tian H, Wu Z, Chen S, Ao K, Huang W, Yaghmaiean H, Sun T, Xu F, Zhang Y, Wang S, et al. 2021. Activation of TIR signalling boosts pattern-triggered immunity. Nature 598(7881): 500–503.

Tsuda K, Mine A, Bethke G, Igarashi D, Botanga CJ, Tsuda Y, Glazebrook J, Sato M, Katagiri F. 2013. Dual regulation of gene expression mediated by extended MAPK activation and salicylic acid contributes to robust innate immunity in Arabidopsis thaliana. PLoS Genet 9(12): e1004015.

Tungadi T, Watt LG, Groen SC, Murphy AM, Du Z, Pate AE, Westwood JH, Fennell TG, Powell G, Carr JP. 2021. Infection of Arabidopsis by cucumber mosaic virus triggers jasmonate-dependent resistance to aphids that relies partly on the pattern-triggered immunity factor BAK1. Mol Plant Pathol 22(9): 1082–1091.

van Wersch S, Tian L, Hoy R, Li X. 2020. Plant NLRs: The Whistleblowers of Plant Immunity. Plant Commun 1(1): 100016.

Wan J, Tanaka K, Zhang XC, Son GH, Brechenmacher L, Nguyen TH, Stacey G. 2012. LYK4, a lysin motif receptor-like kinase, is important for chitin signaling and plant innate immunity in Arabidopsis. Plant Physiol 160(1): 396–406.

Wan WL, Zhang L, Pruitt R, Zaidem M, Brugman R, Ma X, Krol E, Perraki A, Kilian J, Grossmann G, et al. 2019. Comparing Arabidopsis receptor kinase and receptor protein-mediated immune signaling reveals BIK1-dependent differences. New Phytol 221(4): 2080–2095.

Wang G, Ellendorff U, Kemp B, Mansfield JW, Forsyth A, Mitchell K, Bastas K, Liu CM, Woods-Tor A, Zipfel C, et al. 2008. A genome-wide functional investigation into the roles of receptor-like proteins in Arabidopsis. Plant Physiol 147(2): 503–517.

Wang J, Chai J. 2020. Structural Insights into the Plant Immune Receptors PRRs and NLRs. Plant Physiol 182(4): 1566–1581.

Wang Y, Zhang H, Wang P, Zhong H, Liu W, Zhang S, Xiong L, Wu Y, Xia Y. 2023. Arabidopsis EXTRA-LARGE G PROTEIN 1 (XLG1) functions together with XLG2 and XLG3 in PAMP-triggered MAPK activation and immunity. J Integr Plant Biol 65(3): 825–837.

Willmann R, Lajunen HM, Erbs G, Newman MA, Kolb D, Tsuda K, Katagiri F, Fliegmann J, Bono JJ, Cullimore JV, et al. 2011. Arabidopsis lysin-motif proteins LYM1 LYM3 CERK1 mediate bacterial peptidoglycan sensing and immunity to bacterial infection. Proc Natl Acad Sci U S A 108(49): 19824–19829.

Withers J, Dong X. 2017. Post-translational regulation of plant immunity. Curr Opin Plant Biol 38: 124–132.

Wu Y, Gao Y, Zhan Y, Kui H, Liu H, Yan L, Kemmerling B, Zhou JM, He K, Li J. 2020. Loss of the common immune coreceptor BAK1 leads to NLR-dependent cell death. Proc Natl Acad Sci U S A 117(43): 27044–27053.

Yang DH, Hettenhausen C, Baldwin IT, Wu J. 2011. BAK1 regulates the accumulation of jasmonic acid and the levels of trypsin proteinase inhibitors in Nicotiana attenuata’s responses to herbivory. J Exp Bot 62(2): 641–652.

Yeh YH, Chang YH, Huang PY, Huang JB, Zimmerli L. 2015. Enhanced Arabidopsis pattern-triggered immunity by overexpression of cysteine-rich receptor-like kinases. Front Plant Sci 6: 322.

Yip Delormel T, Boudsocq M. 2019. Properties and functions of calcium-dependent protein kinases and their relatives in Arabidopsis thaliana. New Phytol 224(2): 585–604.

Yu TY, Sun MK, Liang LK. 2021. Receptors in the Induction of the Plant Innate Immunity. Mol Plant Microbe Interact 34(6): 587–601.

Yuan M, Cai B, Xin XF. 2023. Plant immune receptor pathways as a united front against pathogens. PLoS Pathog 19(2): e1011106.

Yuan M, Jiang Z, Bi G, Nomura K, Liu M, Wang Y, Cai B, Zhou JM, He SY, Xin XF. 2021a. Pattern-recognition receptors are required for NLR-mediated plant immunity. Nature 592(7852): 105–109.

Yuan M, Ngou BPM, Ding P, Xin XF. 2021b. PTI-ETI crosstalk: an integrative view of plant immunity. Curr Opin Plant Biol 62: 102030.

Zhang W, Fraiture M, Kolb D, Loffelhardt B, Desaki Y, Boutrot FF, Tor M, Zipfel C, Gust AA, Brunner F. 2013. Arabidopsis receptor-like protein30 and receptor-like kinase suppressor of BIR1-1/EVERSHED mediate innate immunity to necrotrophic fungi. Plant Cell 25(10): 4227–4241.

Zhong CL, Zhang C, Liu JZ. 2019. Heterotrimeric G protein signaling in plant immunity. J Exp Bot 70(4): 1109–1118.

Zhou JM, Zhang Y. 2020. Plant Immunity: Danger Perception and Signaling. Cell 181(5): 978–989.

Zhou N, Tootle TL, Glazebrook J. 1999. Arabidopsis PAD3, a gene required for camalexin biosynthesis, encodes a putative cytochrome P450 monooxygenase. Plant Cell 11(12): 2419–2428.

Zhou N, Tootle TL, Tsui F, Klessig DF, Glazebrook J. 1998. PAD4 functions upstream from salicylic acid to control defense responses in Arabidopsis. Plant Cell 10(6): 1021–1030.

